# Modeling the role of urokinase plasminogen activator, uPA, and circulating Cancer-Associated Fibroblasts (cCAFS) in breast cancer cell extravasation

**DOI:** 10.1101/2025.06.11.659108

**Authors:** Angela Spartz, Susan Schmidt, Fazila Mohamed, Benjamin Troness, Carol A. Lange, Dorraya El-Ashry

## Abstract

Circulating Cancer-Associated Fibroblasts (cCAFs) have been discovered in circulating tumor cell clusters from all stages of disease progression in breast cancer patients. We have shown that CAFs promote lung metastases in the mouse tail vein model when they are clustered with triple negative breast cancer (TNBC) MDA-MB231 cells. Following on this observation, we saw that MDA-MB231-luciferase labeled cells persist at higher levels when present in CAF23/MDA-MB231 co-clusters compared to MDA-MB231 mono-clusters within the first 3 days after tail vein injection. This prompted us to investigate whether CAFs aid cancer cell extravasation from capillary venules into the lung parenchyma, which would impart better survival and faster seeding of metastases. Ex vivo lung extravasation assays showed that within the first 8-24 hrs after tail vein injection, more cells from CAF23/MDA-MB231 co-clusters extravasated than cells from MDA-MB231 mono-clusters. Using in vitro endothelial binding assays, we determined that CAF/TNBC co-clusters bind to HUVEC endothelial cells better than TNBC mono-clusters. Single Cell RNA-seq identified several genes in the fibrinolysis pathway whose expression increases in TNBC cells when they are clustered with CAFs. One of these genes is *PLAU*, which encodes the urokinase-type plasminogen activator, uPA. siRNA knockdown of PLAU decreased in vitro TNBC-endothelial cell interactions and ex vivo extravasation of MDA-MB231 mono-clusters, revealing a role for uPA/PLAU in breast cancer cell extravasation. Our data helps to define the role of CAFs in breast cancer extravasation and highlights the importance of our previous work showing that CAFs promote tumor cell dissemination and metastasis.

## Introduction

Metastasis, or the dissemination of tumor cells from a primary site into the circulation to seed new tumors, is the primary cause of death from cancer. A comparison of 5-year survival rates for breast cancer patients with localized disease (99 %) versus distant disease (31%) illuminates the importance of understanding this biological process[1]; if we could slow metastasis, or stop it altogether, cancer occurring in non-essential organs could be nearly curative.

Cancer-associated fibroblasts (CAFs) are the most abundant non-cancerous cell type in the tumor microenvironment (TME). CAFs can evolve from resident fibroblasts or several other cell types, including epithelial cells, pericytes, and bone marrow derived mesenchymal stem cells. CAFs are activated by tumor cell secretion of signaling molecules such as TGFβ, PDGFα, PDGFβ, or FGF2, as well as environmental factors[2, 3]. A characteristic phenotype of CAFs is their hyperactivated state, which includes enhanced secretion of cytokines (e.g. IL6 and IL10), chemokines (e.g. IL8 and CXCL12), and growth factors (e.g. FGF2 and TGFβ).

These signaling molecules impart many of the tumor promoting functions of CAFs. Indeed, CAFs and many of their secreted factors are implicated in nearly all functions defined by the hallmarks of cancer; they promote proliferation, invasion, migration, metastasis, stemness, immunosuppression, altered cancer cell metabolism, and drug resistance [4–6]. This is true for nearly all solid tumor types including breast, ovarian, lung, colorectal, and pancreatic cancer.

We previously isolated and identified circulating CAFs (cCAFs) in clusters with circulating tumor cells (CTCs) from human blood samples at all stages of breast cancer development[7, 8]. Using the PYMT syngeneic model, we correlated a greater abundance of cCAF/CTC co-clusters in mice that presented with late-stage disease[8]. Human primary CAF cell lines can form co-clusters in vitro with human breast cancer cells when grown in ultra-low attachment conditions[8, 9]. The ability of BC cells to co-cluster with CAFs correlates with the abundance of the cell membrane protein, CD44, which is high in triple negative breast cancer (TNBC) cell lines and lower in estrogen receptor (ER) positive breast cancer cell lines[9]; knockdown of CD44 reduces co-clustering of CAFs with either type of breast cancer cell line[8, 9]. Using the tail vein model of metastasis, we showed that CAF23/MDA-MB231 co-clusters made in vitro establish metastases faster than MDA-MB231 monoclusters[8], consistent with previous studies showing that CAFs promote metastasis[10].

One of the most under-studied steps in the metastatic cascade is extravasation (also termed trans-endothelial migration, TEM), the process in which migrating cells exit the microvasculature through a single cell endothelial layer. Increased extravasation can lead to enhanced breast cancer cell survival, accelerated establishment of the metastatic niche, and increased tumor cell proliferation. Early modeling of breast cancer extravasation utilized the rolling model of leukocyte extravasation since both cell types express similar cellular adhesion molecules [11–13]. In this model, cells “roll” along the endothelial layer through E- or P-selectin-mediated, weak cell-cell attachments, which allow cells to move while maintaining physical connections. At the point of exit, cell-cell attachments become stronger through firm adhesions mediated by integrin proteins, and the endothelial cells undergo extracellular matrix (ECM) reorganization. Finally, Rho/ROCK GTPase signaling alters endothelial cell permeabilization through the disassembly of adherens junctions and changes in the actin cytoskeleton[12]. In a process known as diapedesis, extravasating cells break through (transcellular migration) or between (paracellular migration) endothelial cells and transmigrate out of the circulation.

Various studies have shown that some cancer cells follow the rolling model of extravasation; however, there also appears to be some variation. For instance, CTCs and CTC clusters (which are larger than leukocytes) often become trapped in small diameter vessels due to physical restriction; although evidence for cancer cell arrest in wider vessels through the formation of strong adhesions also exists[14]. While diapedesis is predominant in leukocyte TEM, larger CTCs sometimes do not make it across the capillary wall without causing severe damage or even death to endothelial cells. Additionally, the cellular adhesion and signaling molecules that are involved at each step of extravasation may differ between leukocytes and CTCs and even among different types of cancer cells.

Secreted factors, such as VEGF[15], ANGPT2, ANGPTL4[16] and CCL2[17], play important roles during extravasation as they can alter the expression of cellular adhesion molecules and activate signaling pathways (such as integrin and Rho/ROCK signaling) important for regulating vascular permeability and ECM reorganization [18]. Secreted factors can originate from CTCs themselves[17] or from other cells, such as endothelial cells, platelets, or macrophages, and certainly from CAFs[9]. On the other hand, CTCs or cells interacting with them (linker cells, such as platelets) may directly bind with endothelial cells through adhesion molecules on their surfaces. A subset of CAFs, termed matrix CAFs or mCAFs, express abundant adhesion molecules and ECM proteins[19, 20].

Since cCAF/CTC clusters have been identified in breast cancer patients and CAFs have a profound effect on the behavior of cancer cells, we hypothesized that CAFs may enhance breast cancer cell extravasation. Here, we have investigated the influence of CAFS on extravasation using in vitro binding assays and ex vivo lung extravasation assays. Additionally, we have performed Single Cell RNA sequencing (scRNA-seq) to investigate the cell-cell communication that occurs between CAFs and breast cancer cells under in vitro, non-adherent (3D) conditions. We identify the plasminogen urokinase, or uPA/PLAU, as a CAF23 upregulated gene, and show that it is an important molecular player in the process of breast cancer cell extravasation. Our data demonstrate that CAFs aid breast cancer cell extravasation and underscore the significance of the dissemination of CAFs with breast cancer cells, a process that likely occurs in other disseminating solid tumors as well.

## Materials and Methods

### Cell culture

MDA-MB231 and DT28 cell lines were cultured in IMEM (Gibco)+ 10% Fetal Bovine Serum (FBS) (GE Healthcare). MDA-MB468 was cultured in DMEM/F12 50:50 (Corning) with 10% FBS and 1% penicillin-streptomycin (Gibco). SUM159 was cultured in DMEM/F12 50:50 with 10% FBS, 1 μg/ml insulin (Gibco), 1 μg/ml hydrocortisone (StemCell Technologies) and 1% penicillin-streptomycin. Primary CAF19 and CAF23 cells (passage 15-20) were cultured in gelatin-coated (Cell Biologics) flasks in IMEM + 20% FBS. The DT28 primary TNBC cell line and primary human CAF cell lines, CAF19 (RRID:CVCL_D4YH) and CAF23 (RRID:CVCL_D4YJ), were derived from primary breast tumor specimens and have been previously characterized [21]. HUVEC cells were purchased from Sigma Aldrich (cat # 200P-05N) and cultured in gelatin-coated flasks with Endothelial Cell Growth Medium (Cell Applications); for experiments, HUVEC cells were used between passages 2-6. Dextran-coated, charcoal-stripped FBS (Hyclone) was used in place of FBS for experiments where starvation was required before TGFβ administration. Cells were tested monthly for mycoplasma with the Mycoalert Mycoplasma detection kit (Lonza) or by eMyco Plus PCR kit (Lilif Diagnostics). All cells were maintained under 5% CO in a forced air incubator at 37°C as per standard tissue culture practices. For tail vein metastasis studies, breast cancer cells were transduced with pLenti CMV Puro LUC (w168-1) (Addgene plasmid # 17477) and maintained with 0.5 μg/ml puromycin (VWR). Luciferase activity was assessed using the Luciferase Assay System (Promega).

### Chemicals

The following chemicals were used in our studies: Bovine Hyaluronidase (StemCell Technologies), 4MU (Sigma Aldrich), UK122 (MedChemExpress), TGFβ1 (R and D Systems), Cell Tracker Deep Red (Invitrogen, C34565) or Cell Tracker Red (Invitrogen, C34552), FITC-lectin (Vector Laboratories). The following antibodies were used for western blots: anti-PLAU (ProteinTech 17968-1-AP), anti-CD44-AF647 (Novus Biologicals, MEM-263), biotinylated HA binding protein (HABP) (EMD Millipore), AF647-Streptavidin (BioLegend 405237), horseradish peroxidase(HRP)-conjugated goat anti-mouse (Biorad), and horseradish peroxidase (HRP)-conjugated goat anti-rabbit HRP (Biorad).

### siRNA and shRNA knockdowns

siRNA knockdown of CD44 was performed 3 days prior to experiments as described [8] using 20 nM CD44 siRNA. siRNA knockdown of PLAU was performed for 3 days prior to experiments using 20 nM siPLAU and 3 μl Silentfect (Biorad) in 3 ml final volume growth media according to manufacturer’s instructions. All siRNAs were purchased from Dharmacon with the following catalog numbers: non-targeting control (L-001810-10-05), CD44 (L-009999-00-00-0005), PLAU (L-00600-00-0005).[9]

### Real-time PCR

Real-time PCR was performed as previously described[9] with the modification that RNA was treated with DNAse1 (Qiagen) according to manufacturer’s instructions prior to cDNA formation. PCR primers used include the following: PLAU: (F: 5’TGCCCTCGATGTATAACGATCC, R: 5’ GGTGGGAAATCAGCTTCACAAC) 18S rRNA: (F: 5’GGAGAGGGAGCCTGAGAAAC, R: 5’ TCGGGAGTGGGTAATTTGC).

### Immunofluorescence

Cells for immunofluorescence were grown on coverslips in 6-well dishes, washed with PBS, fixed with 4% paraformaldehyde for 20 minutes, and then permeabilized in PBS + 0.1% Triton X-100 for 20 minutes. After 2 more washes, coverslips were blocked for 1 hr at room temperature with blocking buffer (3% normal goat serum, 1% bovine serum albumin in 1x PBS). Coverslips were incubated with 1:1000 anti-CD44-647 overnight at 4°C. The next day, coverslips were washed several times in PBS, and mounted onto slides with Prolong Gold anti-fade plus DAPI. For hyaluronin staining, cells were fixed and permeabilized as above, then blocked in blocking buffer (as above without normal goat serum) for 1 hr at room temp. Coverslips were then incubated with 1:100 HABP for 1 hr, washed, and incubated with 1:500 Alexafluor647-streptavidin for 30 min at 4°C. After several additional washes, coverslips were mounted onto slides with Prolong gold anti-fade plus DAPI (Invitrogen).

### Western blots

Cells were harvested and lysed in radioimmunoprecipitation assay lite (RIPA-lite) buffer as previously described[9]. Western blots were blocked with 5% non-fat dry milk at room temp for 1 hr and incubated with anti-PLAU antibody (1:1000) at 4°C overnight or anti-GAPDH (1:1000) at room temp for 1 hr. After washing, blots were incubated with 1:5000 HRP-conjugated secondary antibodies for 1 hr and processed for chemiluminescence detection.

### In vitro clusters and in vitro clustering assay

To make co-clusters or mono-clusters, BC cells and CAF cells were dissociated using the Accutase (Gibco) gentle dissociation reagent and made into a single cell suspension. Equal numbers of BC cells and CAF cells were either plated alone or together into ultra-low-attachment plates in Mammocult medium (StemCell Technologies) according to the manufacturer’s instructions and incubated at 37°C and 5% CO_2_ for 24 or 48 hours. Quantitative in vitro clustering assays were performed as described[9] using 5,000 cells of each type.

### Endothelial binding assay

HUVEC cells were seeded into a 12-well plate and allowed to grow for 3 days to form a confluent monolayer. Media was refreshed on Day 2. On the day of the binding assay, breast cancer cells and CAF cells were labeled with Cell Tracker dyes for 30 minutes at 37°C in PBS and allowed to recover in regular culture media for 2 hours. BC and CAF cells were lifted with Accutase and resuspended in IMEM and counted to achieve a final cell concentration of 2,000 cells/sample well. Endothelial cell media was carefully removed from the HUVEC monolayer, replaced with 1 ml of cells (in IMEM), and returned to the incubator for 30 min. At the end of the assay, unbound cells were carefully but quickly washed 2X with fresh IMEM. Plates were scanned on a Nikon TiE microscope at the appropriate wavelengths to detect Cell Tracker dyes. To obtain accurate counts of percent bound cells, 3 control wells containing the same concentration of starting cells were made, allowed to adhere, scanned and their numbers averaged. Thus, percentage of bound cells was calculated as # cells bound/# starting cells. In experiments where clusters were used, 15-50K cells of each labeled cell type were used to make 24-hr clusters. Clusters were pooled, resuspended in an appropriate volume of IMEM media, and evenly aliquoted to sample wells or control wells. Only clusters with 5 or more cells were counted for analysis. For experiments using CAF conditioned media, CAF cells were grown in regular IMEM + 10% FBS until they were 75% confluent, washed 2X with PBS and then 10 mls IMEM (no phenol red) was added. Cells were grown for 2 additional days, media was collected, and filtered through a 0.2 μM syringe filter (PALL Life Sciences). For the hyaluronin inhibition experiment, cells were treated overnight with 1 mM 4-methyl-umbelliferone (4-MU). The next day cells were stained with Cell Tracker dyes for 30 min, lifted with Accutase, and treated with 2000 units/ml of bovine hyaluronidase for 1 hr at 37°C. Cells were then counted and used directly in the endothelial binding assay.

### Tail vein metastasis model

75,000 MDA-MB231-Ffluc cells and or 75,000 MDA-MG231-Ffluc + 75,000 CAF23 cells per well per mouse were allowed to form in vitro clusters as above for 24 hrs. Clusters were pooled and resuspended in PBS to a volume of 100ul/mouse. Injections were done via tail vein into 6-week-old female NOD.Cg-*Prk-dc^scid^ Il2rg^tm1Wjl^*/SzJ (NSG) mice (Jackson Laboratories, cat# 005557) without anesthesia, and animals were returned to housing facilities until the time of imaging. For luminescence imaging, mice were injected i.p. with 100 μl of IVISbrite D-luciferin Ultra Rediject Substrate (Revvity) 10 minutes prior to isoflurane anesthesia and imaging. Imaging was done on a Bruker Extreme Optical Small Animal Imaging System (Bruker) at 2 hrs, 18 hrs, 3 days and 5 days after tail vein injection of cells and were further imaged every week thereafter to track tumor progression. Analysis of luminescence was done using Bruker Imaging Analysis software. Animal protocols for this study were approved by the University of Minnesota Institutional Animal Care and Use Committee (IACUC). Animals were housed 5 per cage in U of MN Research Animal Resources (RAR) facilities and allowed to acclimate for one week prior to handling.

### Ex vivo extravasation assays

BC cells and CAFs were labeled with Cell Tracker Red or Cell Tracker Deep Red respectively, for half an hour at 37°C and then allowed to recover for 2 hrs in their regular culture media. Cells were lifted and made into in vitro clusters as above using 75,000 BC + 75,000 CAF (for co-clusters) or 150,000 BC cells (for mono-clusters) per well of a 6-well ULA plate. At 24 hrs, 2 wells of clusters were pooled and resuspended into 100 μl PBS for injection per mouse. Six-week-old female FVB/Nj (Jackson Laboratories, cat# 001800) mice were injected with clusters via tail vein and returned to animal facilities until the specified time point (8 hrs 24 hrs or 48 hrs). One hour before lung collection, each mouse was injected by tail vein with 100 μl FITC-tomato lectin (Vector Labs) diluted 1:2 in PBS. For lung collection, the mouse was anesthetized, and cardiac puncture was performed to remove most of the blood, the chest was opened, and lungs were perfused through the trachea with 2% low melt agarose. Lung tissue was placed in PBS + 1% formalin for 1 hr and imaged immediately or stored in PBS for up to 7 days. Lung tissue was sliced thinly with a razor blade and arranged in a glass bottom culture dish (Ibidi). Z-stack images from 10-20 clusters/mouse were taken on a Nikon A1R or Nikon AXR confocal microscope and evaluated for extravasation. A cluster was considered partially extravasated if either the CAF cells or BC cells could be observed partially outside of the vasculature.

### Single Cell RNA sequencing

48-hr in vitro mono-clusters and co-clusters were made for Single Cell RNA sequencing as above using the following combinations of cells: MDA-MB231, CAF23 + MDA-MB231, DT28, CAF23 + DT28, and CAF23. 240,000 cells were used for each sample and samples with 2 types of cells were used at a 1:1 ratio. To remove any cells in the dishes that had not formed part of a cluster, the cells were filtered over a 15 µm Pluristrainer (Pluriselect 43-50015-50) and then inverted to release the clusters. Clusters were then digested with Accutase to create a single cell suspension. Cells were washed with PBS in LoBind tubes (Eppendorf) and resuspended in PBS. Single cell preparations (grown under adherent conditions) were also made for each cell line. Each sample was incubated with a separate Hashtag Oligo antibody (Biolegend, Supplemental Table 1); two different experiments were performed at different times and therefore HTO10 was used for two different samples. Stained cells from each sample were given to the University of Minnesota Genomics Center to be made into 10x Chromium 3’GEX cDNA libraries (10X Genomics). For in depth sequencing, libraries were pooled together and run on a single Novaseq gel (Illumina).

### ScRNA-seq Analysis

CellRanger (10x Genomics) was used to perform counting of each barcoded unique molecular identifier (UMI) and the resulting raw feature matrix files were analyzed with Seurat (V4) [22]. For experiments 1 and 2, sequencing saturation was 37.9% and 36.8% respectively. To analyze the quality of the sequencing and mapping, we evaluated the percentage of reads that mapped to the transcriptome, exons, introns, and intergenic regions and considered them to be within acceptable ranges. Median UMI counts/cell was about 11,000 and median genes/cell was about 3,000 for each experiment, with 25,000-30,000 cells total for each experiment. Preprocessing of the UMIs included reducing the number of unique features to a range of 750-6000 and a percentage of mitochondrial DNA reads to less than 5-10%. Several permutations of these two parameters were tested to obtain the most robust differences in differentially expressed genes (p value < 0.005 with the highest avg_Log2FC) for each comparison made. Demultiplexing was performed with a positive quantile of 0.95-0.99, the quality of Hashtag Oligo (HTO) staining was assessed, and doublets (cells which stained for 2 HTOs) and negatively stained cells were removed. Clustering was performed with the FindClusters command feature in Seurat with a resolution of 0.4-0.6, and 30 dimensions; for ease of analysis, we adjusted the resolution to produce less than 10 clusters per sample. Principle component analysis and non-linear dimensional reduction (UMAP) was run with 30 dimensions. For differential expression analysis, we performed several different algorithms including roc, MAST, and DESeq2. Number of cells analyzed for each graph is indicated in its corresponding figure legend. For IPA (Qiagen) analysis of BC cells, we used an avg_Log2FC cutoff of ±0.25 and adjusted p value of 0.05 to obtain a list of at least 100 DEG for analysis. For IPA analysis of CAF cells, we used an avg_Log2FC cutoff of ±0.5 and adjusted p value of 0.05 to obtain a list of DEG for analysis (112 for DT28 comparisons and 300 for MDA-MB231 comparisons). We also used the fgsea (v 1.32.2) package to analyze CAF single cells compared to MDA-MB231 single cells with Gene Ontology biological function gene sets and the Gene Ontology Resource (geneontology.org) with the Cellular Component gene sets to compare breast cancers cells from mono-clusters vs co-clusters. All graphs were made using Seurat(v4) and ggplot2 (v 3.5.1) in RStudio (v 4.4.0).

## Results

### CAFs aid the early establishment of MDA-MB231 metastases

We have previously reported that CAF23/MDA-MB231 co-clusters injected into the mouse tail vein establish lung metastases faster than MDA-MB231 mono-clusters[8]. We decided to take a closer look at the timing of tumor growth in this model by following the luminescent signal of MDA-MB231-luciferase labeled clusters at multiple time points within the first week after injection. As before, MDA-MB231-luciferase breast cancer cells were allowed to form in vitro clusters in the presence or absence of unlabeled CAF23 cells for 24 hrs. These clusters were injected into the tail vein of immunosuppressed NSG mice and luminescence was imaged at 2 hrs, 18 hrs, 3 days and 5 days after injection. As shown in Figure 1, both types of clusters produced similar luminescence readings at 18 hrs post-injection (relative to the starting reading at 2 hrs post-injection). However, luminescence from mono-clusters decreased more than luminescence from co-clusters on day 3. On day 5, luminescence from co-clusters recovered to the 18-hr level whereas luminescence from mono-clusters did not recover as much. These data suggest that CAFs support the survival and the proliferation of breast cancer cells in the tail vein assay.

**Figure 1.**
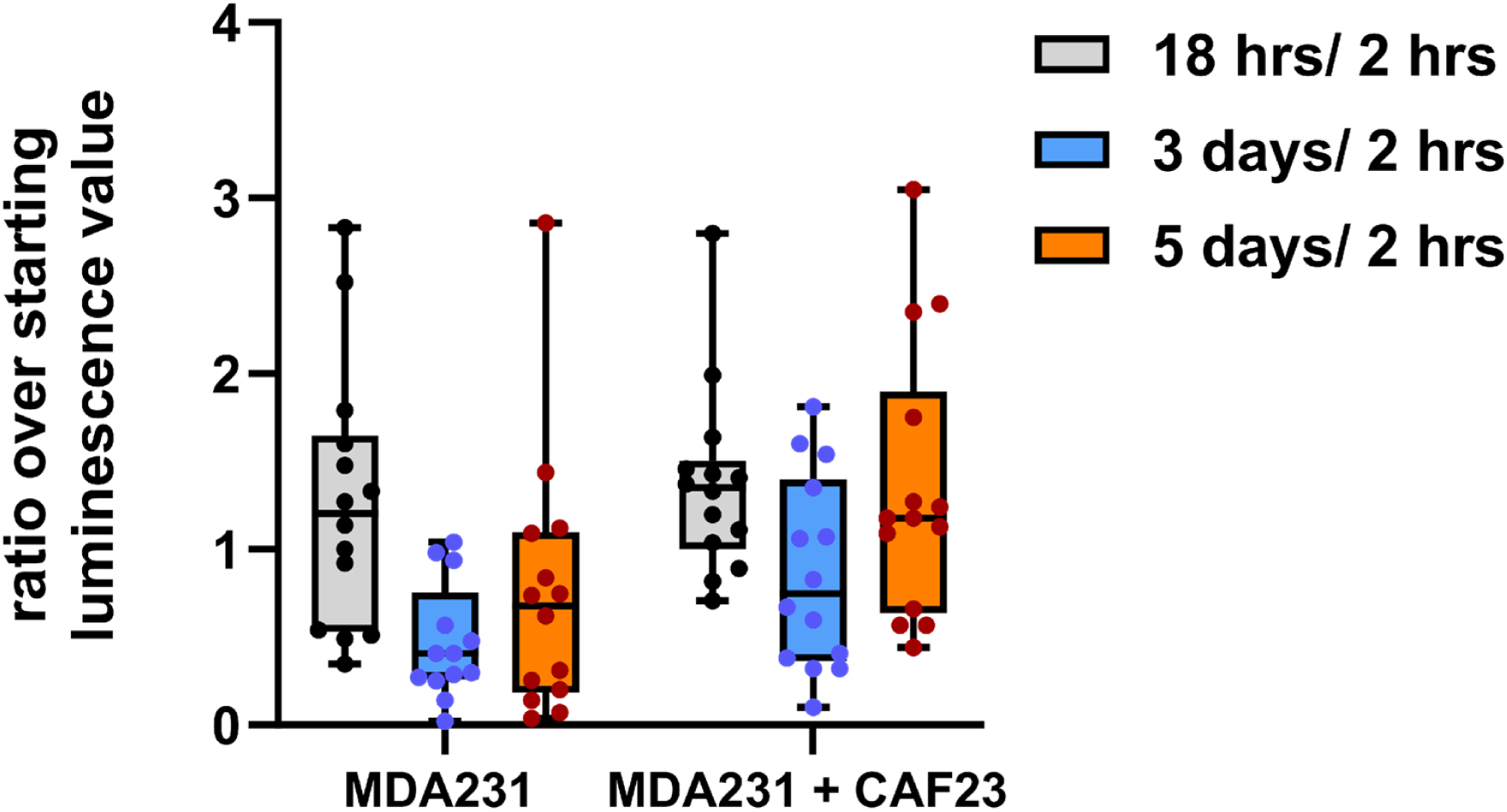
CAF23/MDA-MB231 co-clusters survive and grow faster than MDA-MB231 mono-clusters in the tail vein assay. MDA-MB231-luciferase mono-clusters and CAF23/ MDA-MB231-luciferase co-clusters were made for 24 hrs in Mammocult media (75,000 cells of each cell type per mouse). Clusters were collected into PBS, injected into the tail vein of NSG mice, and luminescence was followed over the first 5 days using live animal imaging. The luminescence observed at each time point (18 hrs, 3 days, and 5 days) was normalized to the amount that was present 2 hrs after injection. Each dot represents a single animal. N = 14 mice per group.

### CAF23/MDA-MB231 co-clusters extravasate faster than MDA-MB231 mono-clusters

The circulatory system is a hostile environment, exhibiting high shear stress forces in venules and capillaries that could contribute to cell death of injected MDA-MB231 cells. If CAFs increase the rate of extravasation, this may increase viability and, ultimately, faster establishment of metastases. To investigate this possibility, we used an ex vivo approach to monitor extravasation of cells from lung capillaries (Fig. 2A). MDA-MB231 and CAF23 cells were labeled with Cell Tracker Red or Cell Tracker Deep Red respectively before making 24-hr in vitro mono-clusters or co-clusters. The clusters were then injected into the tail vein of FVB mice. Half an hour before designated time points (8 hrs, 24 hrs, 48 hrs), FITC-labeled lectin (an endothelial cell binding protein) was injected into the tail vein to visualize the vasculature. Lungs were perfused and extracted, cut into thin sections, and clusters were scored for extravasation by confocal microscopy. Within the venules, clusters had broken apart, but CAFs and cancer cells appeared to remain in physical contact. Usually, only a portion of the cells within a cluster had extravasated; rarely did we see an entire cluster of cells completely extravasated by 48 hrs. We observed that at 8 hrs and 24 hrs significantly more CAF23 and MDA-MB231 cells from co-clusters were partially extravasated than MDA-MB231 cells from mono-clusters (Fig. 2B, C). By 48 hrs, the difference was less pronounced, but still significant.

**Figure 2.**
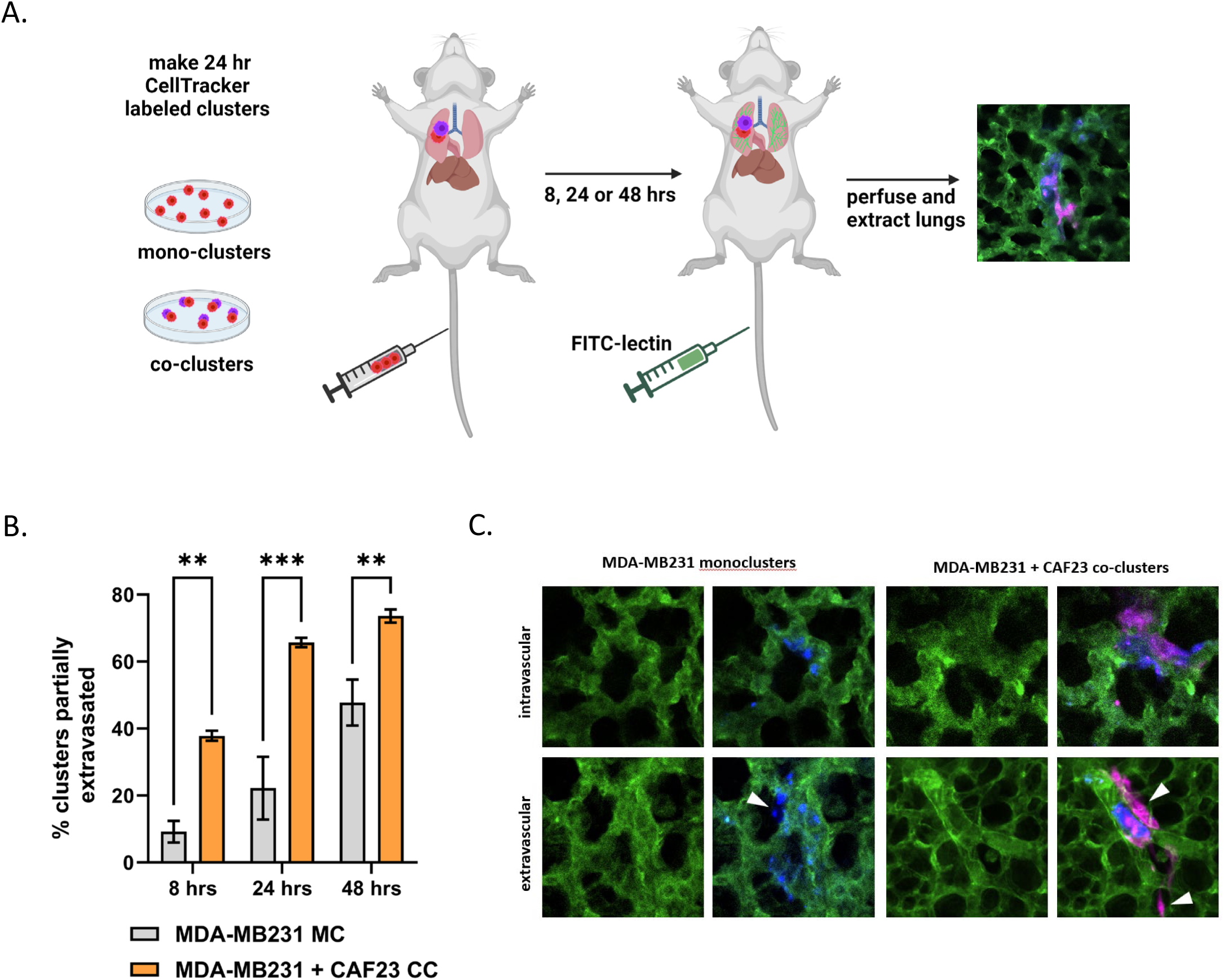
CAF23/MDA-MB231 co-clusters extravasate better than MDA-MB231 mono-clusters in the ex vivo lung extravasation assay. A. Schematic of experimental design. CAF23 cells were labeled with Cell Tracker Deep Red and MDA-MB231 cells were labeled with CellTracker Red. MDA-MB231 mono-clusters and CAF23/MDA-MB231 co-clusters were made for 24 hrs. Clusters were collected and injected into the tail vein of FVB mice. 30 minutes prior to perfusion, FITC-lectin was injected into the tail vein to visualize the vasculature. Lung perfusion and fixation was performed at 8 hrs, 24 hrs, and 48 hrs. Pieces of lung were cut and imaged with confocal microscopy to visualize and quantify cell clusters inside or outside of the vasculature. B. Quantitation of ex vivo assay using 3 mice per time point per treatment. At least 20 clusters per animal were counted and scored as being intravascular or partially extravascular. C. Example images of mono-clusters or co-clusters that are intravascular or partially extravascular. Vasculature is green. MDA-MB231 cells are pseudo-colored blue. CAFs are pink. Arrowheads indicate cells that have extravasated. ** p ≥ 0.005, *** p ≥ 0.0005

### In vitro endothelial binding assays demonstrate enhanced binding with CAF23 cells or CAF23 conditioned media

Physical interactions among CAFs, breast cancer cells, and endothelial cells must occur before extravasation takes place. Models of CTC extravasation maintain that strong cellular interactions precede extra cellular matrix reorganization and trans-endothelial migration. We therefore used an in vitro endothelial binding assay to investigate the cellular interactions among breast cancer cells, CAFs, and endothelial cells. MDA-MB231 and DT28 TNBC cells were used for these assays. Cell Tracker-labeled TNBC mono-clusters or CAF23/TNBC co-clusters were made in vitro for 24 hrs and then equally aliquoted to wells containing a confluent lawn of HUVEC endothelial cells. Clusters were allowed to bind for 30 minutes, and then unbound clusters were gently washed away and the remaining fluorescent clusters were counted. We observed that CAF23/TNBC co-clusters bound to endothelial cells better than TNBC mono-clusters (Fig. 3A). To determine if there was a difference in binding for different primary CAF cell lines, we compared the indolent CAF19 cell line isolated from a luminal A human breast cancer with the CAF23 cell line that was isolated from an aggressive TNBC. CAF mono-clusters made with either CAF19 or CAF23 cell lines bound to endothelial cells with similar efficiency (Fig. 3B). Endothelial binding assays were performed with single cells rather than clusters to determine how well individual TNBC cells bound to endothelial cells in comparison to CAFs. We observed that both CAF cell lines bound better than four different TNBC cell lines (MDA-MB231, DT29, SUM159 and MDA-MB468) (Fig. 3C). In comparison with each other, the TNBC cell lines bound with the same efficiency with the exception of MDA-MB468, which was only slightly less efficient.

**Figure 3.**
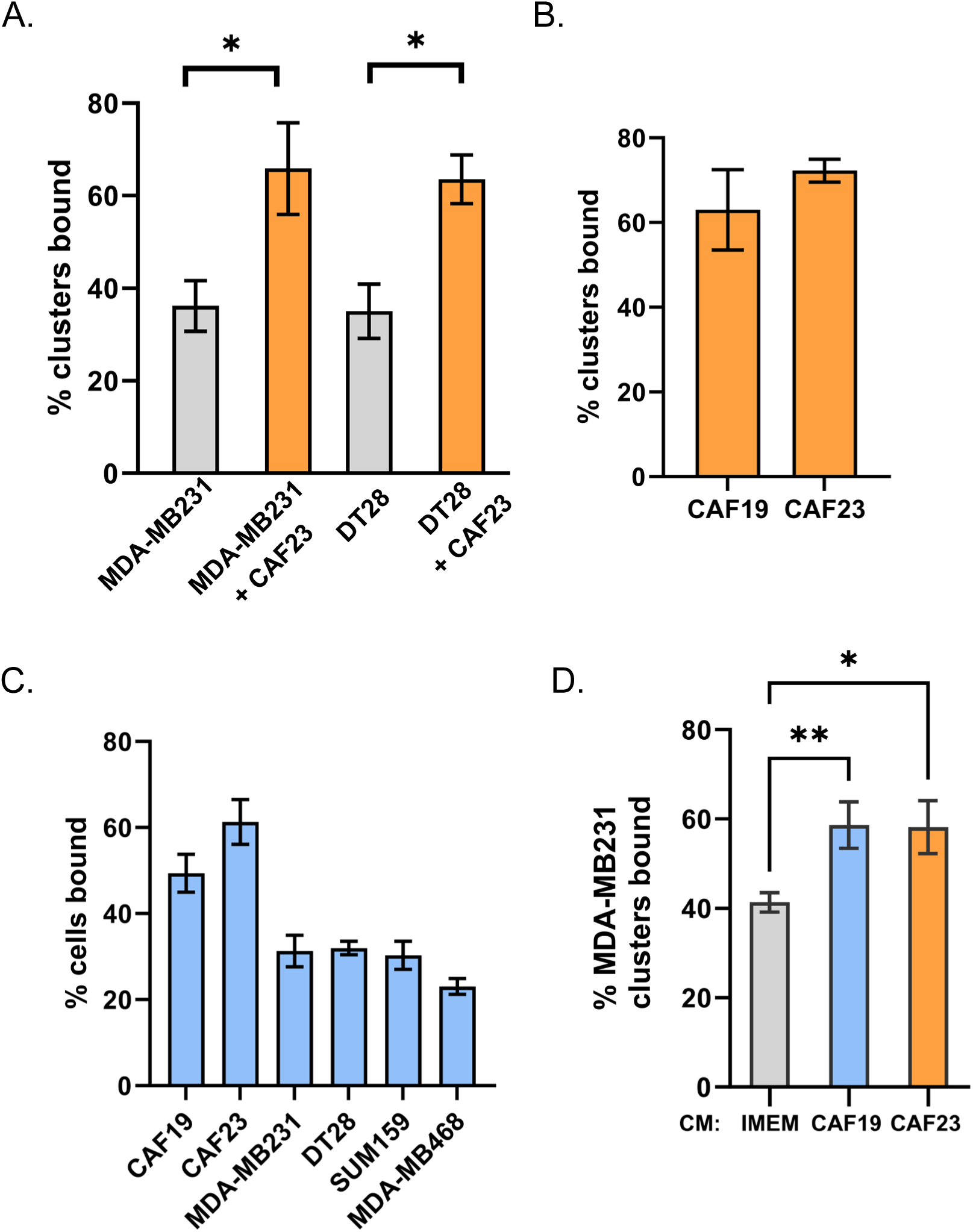
CAFs and CAF/BC co-clusters bind to HUVEC endothelial cells better than BC cells and BC mono-clusters. Assays were performed with CellTracker Deep Red labeled CAFs and CellTracker Red labeled breast cancer cells. Single cell suspensions or 24-hr clusters were made and pipetted over a lawn of HUVEC cells for 30 minutes before being washed. Wells were scanned on a TiE fluorescent microscope and bound cells or clusters were counted. A. Comparison of co-clusters and mono-clusters. B. Comparison of CAF19 and CAF23 mono-clusters. C. CAF cells compared with 4 different TNBC cells. D. Comparison of MDA-MB231 mono-clusters made in CAF conditioned media vs IMEM media. The endothelial binding assay was performed in IMEM. * p ≥ 0.05, ** p ≥ 0.005

Circulating CAF/BC clusters isolated from the blood of breast cancer patients display a random arrangement of CAFs and breast cancer cells with some CAFs on the outside as well as on the inside of clusters[8]. However, clusters made in vitro tend to arrange themselves with the CAFs on the inside with breast cancer cells surrounding them. In our in vitro endothelial binding assays, co-clusters bind better than mono-clusters even though CAFS may not be directly contacting endothelial cells. This suggests that CAFS may alter the protein composition of breast cancer cell membranes or the breast cancer extracellular matrix (ECM) to increase attachment to endothelial cells. CAFS secrete extracellular vesicles containing adhesion proteins that may be transferred to breast cancer cells; evidence for this has been described[23]. Alternatively, CAFs secrete high amounts of paracrine factors such as such as TGFβ, FGF2, SDF1/CXCL12, IL6, and IL8[9] that may ultimately change the gene and protein expression of adhesion proteins in associated cancer cells. To test whether direct cell contact is necessary for CAFs to increase endothelial binding, we performed the assay using CAF conditioned media (CM). In this assay, MDA-MB231 cells were starved in 1% charcoal-stripped FBS (DCC) under adherent conditions for 24 hrs to “reset” the cells. DCC was used to eliminate the effects of hormones. Then cells were labeled and mono-clusters were made in the presence of CAF CM +1% DCC or IMEM +1% DCC. After 24 hrs, clusters were evaluated in the endothelial binding assay. We observed that when mono-clusters were made in either CAF19 or CAF23 CM, the clusters bound better relative to control media (Fig. 3D). This suggests that direct contact with CAFs is not necessary for CAFs to influence BC-endothelial cell binding.

### Hyaluronin present on CAFs may mediate CAF binding to endothelial cells but CD44 does not

The adhesion protein CD44 and its binding partner, hyaluronin (HA), are moderately expressed on MDA-MB231 cells and highly expressed on CAFs (Supplemental Fig. 1A, C left side, Supplemental Fig. 2C). CD44 and HA mediate clustering between CAFs and MDA-MB231 TNBC [8]. Extracellular vesicles secreted from CAFs contain CD44[23]. CD44 is also reported to play a role in leukocyte rolling and extravasation[18]. We therefore decided to test whether CD44 mediates endothelial binding of CAFs and breast cancer cells. siRNA of CD44 in either CAF23 or MDA-MB231 cells effectively decreased CD44 levels (Supplemental Fig. 1A); however, it did not affect their ability to bind to HUVEC endothelial cells (Supplemental Fig. 1B). Decreasing HA by combination treatment with 4MU overnight and hyaluronidase for 1 hour visibly decreased HA in both cell types (Supplemental Fig. 1C) and decreased binding of CAF23 cells to endothelial cells (Supplemental Fig. 1D). This treatment did not have a significant effect on MDA-MB231 binding to endothelial cells. These data suggest that HA contributes to efficient CAF-endothelial binding, though whether this is through CD44 expressed on HUVEC cells remains to be determined.

### Single Cell RNA-seq reveals cross-talk between CAF23 and TNBC cells

Cross-talk between CAFs and breast cancer cells when they are in cCAF/CTC clusters has not been investigated, and such studies are difficult to accomplish due to the low numbers of circulating CAF/BC co-clusters observed in patients (the presence of cell clusters is lower than that of rare CTCs). We therefore decided to investigate the cross-talk between our CAF23 cell line and BC cells when they are in contact using in vitro co-clusters coupled with the technology of Single Cell RNA-seq (scRNA-seq). Several samples were made and compared for our analysis: 1) MDA-MB231, CAF23, and DT28 single cells grown in 2D with their standard media, 2) CAF23 mono-clusters, MDA-MB231 mono-clusters, and DT28 mono-clusters grown in 3D with Mammocult media for 48 hrs, and 3) CAF23/MDA-MBA231 co-clusters and CAF23/DT28 co-clusters grown in 3D with Mammocult media for 48 hrs. Since not all cells in the clustering assay became part of a cluster, we used Pluriselect filters to remove non-clustered cells, and then used Accutase to disrupt the clusters. A single cell suspension of each sample was labeled with Hashtag Oligos (HTOs) (Supplemental Table 1), and cDNA libraries were made with the 10xGenomics platform. For in silico analysis, we used 10XGenomics Cell Ranger to generate counts and Seurat (v4) for downstream analysis.

First, we sought to characterize the CAF23 cell line since this had not been done previously on a single cell level. CAF23 cells were originally isolated from a dissociated human TNBC tumor sample that by microarray analysis clustered with mammoplasty fibroblasts and mesenchymal stem cells rather than their tumor of origin. Subsequently, these cells were confirmed to be CAFs by the expression of Fibroblast Activation Protein (FAP), Vimentin, and smooth muscle actin (SMA)[21]. ScRNA-seq analysis of CAF23 single cells grown in 2D confirmed the expression of several markers that are commonly used in the literature to identify CAFs, such as VIM, PDGFRa/b, ACTA2, and FAP (Supplemental Fig. 2). The majority of these markers were evenly distributed throughout the CAF23 population, but we were surprised to find that ACTA2 (smooth muscle actin) was present only in a subpopulation of CAFs. ACTA2 is often used as a marker to identify CAFs, but it is more highly expressed on CAFs of vascular origin[20].

We then compared our CAF23 single cell population to MDA-MB231 single cells in order to identify a set of CAF-specific genes that are not expressed by cancer cells for use in subsequent analysis (Supplemental Fig. 3A, B, C). First, using our HTOs, we separated the CAF23 cells from MDA-MB-231 cells (Supplemental Fig. 3A). Then we found a set of markers that were upregulated in CAF23 cells while showing little to no expression in MDA-MB231 cells (Supplemental Fig. 3B). Top genes identified included: FAP, COL5A3, COL6A2, VCAN, DDR2, and EVAB1 (Supplemental Fig. 3B, C). Other researchers have identified sub-categorizations of CAFs isolated from human cancers[20, 24]; these studies have separated CAFs into anywhere from 4-8 sub-categorizations that appear to align with their cell of origin, such as mCAFs (matrix CAFs), vCAFs (vascular CAFs), and iCAFs (inflammatory CAFs). We performed Gene Set Enrichment Analysis with the Gene Ontology Cellular Component gene sets and found several gene sets that are upregulated in CAF23 cells compared to MDA231 cells, such as collagen trimers, extra cellular matrix, extra cellular space, and cell junctions (Supplemental Fig. 3D). These data suggest that our CAF23 cell line most closely resembles mCAFs (matrix CAFs). Consistent with this, our CAF23 cell line expressed genes identified in other studies as highly expressed in mCAFs, including COL11A1, VCAN, and FN1[19, 20]. Known genes from the other CAF sub-classifications were not highly expressed in our CAF23 cell line, such as: 1) CFD and PLA2G2A (iCAFs), 2) PRG4, MYH11, MCAM and RGS5 (vCAFs), 3) NDRG1, PDPN, and MME (tumor-like CAFs or tCAFs). For the tCAF sub-classification, we did see high expression of the stress response genes, ENO1 and GAPDH, but none of the other genes associated with that sub-class.

We next sought to determine what genes are upregulated in TNBC cells by CAFs when they are in co-clusters for 48 hrs. HTOs identified MDA-MB231 mono-clusters from CAF23/MDA-MB231 co-clusters, as expected (Supplemental Fig. 4A). Nearest neighbor clustering and UMAP analysis revealed a group of cells expressing CAF markers (VCAN, PDGFa, PDGFb, FAP, COL6A3, and COL5A2) clustered separately from other cells; these cells were presumed to be CAF23 cells from the co-cluster sample (Supplemental Fig. 4A). Surprisingly, BC cells from mono-cluster and co-cluster samples did not separate from each other on a UMAP plot, suggesting that being in a co-cluster with CAF cells only slightly changed the transcriptional profile. Similar analysis was performed with DT28 mono-clusters and co-clusters with similar results (Supplemental Fig. 5A). We removed the CAF cells from the analysis based on their CAF markers and redid normalization, principal component analysis, nearest neighbor, and UMAP analysis. We then performed differential gene expression analysis, comparing BC cells coming from mono-clusters versus BC cells coming from co-clusters, (474 mono-cluster cells vs 902 co-cluster cells). Overall, average Log2xFC change values were low (−1.2 – 0.85); of the 103 genes that had a Log2xFC of at least ±0.25, 58 were upregulated and 45 were downregulated in the presence of CAFs. While this indicated a subtle difference in the average gene expression values, we observed a more obvious difference in the percentage of cells that exhibited upregulation or downregulation of a given gene (Supplemental Fig. 4B). This can also be seen in a heatmap of the top 10 DEG across all BC cells from each sample (Fig. 4A). In general, it appeared that being in a CAF co-cluster did not result in gene expression changes across all cells in the population but rather in just some BC cells. This may be expected since some BC cells were physically closer to CAFs than others. Analysis of BC cells from DT28 mono-clusters versus CAF23/DT28 co-clusters revealed similar trends (Supplemental Fig. 5B, C).

**Figure 4:**
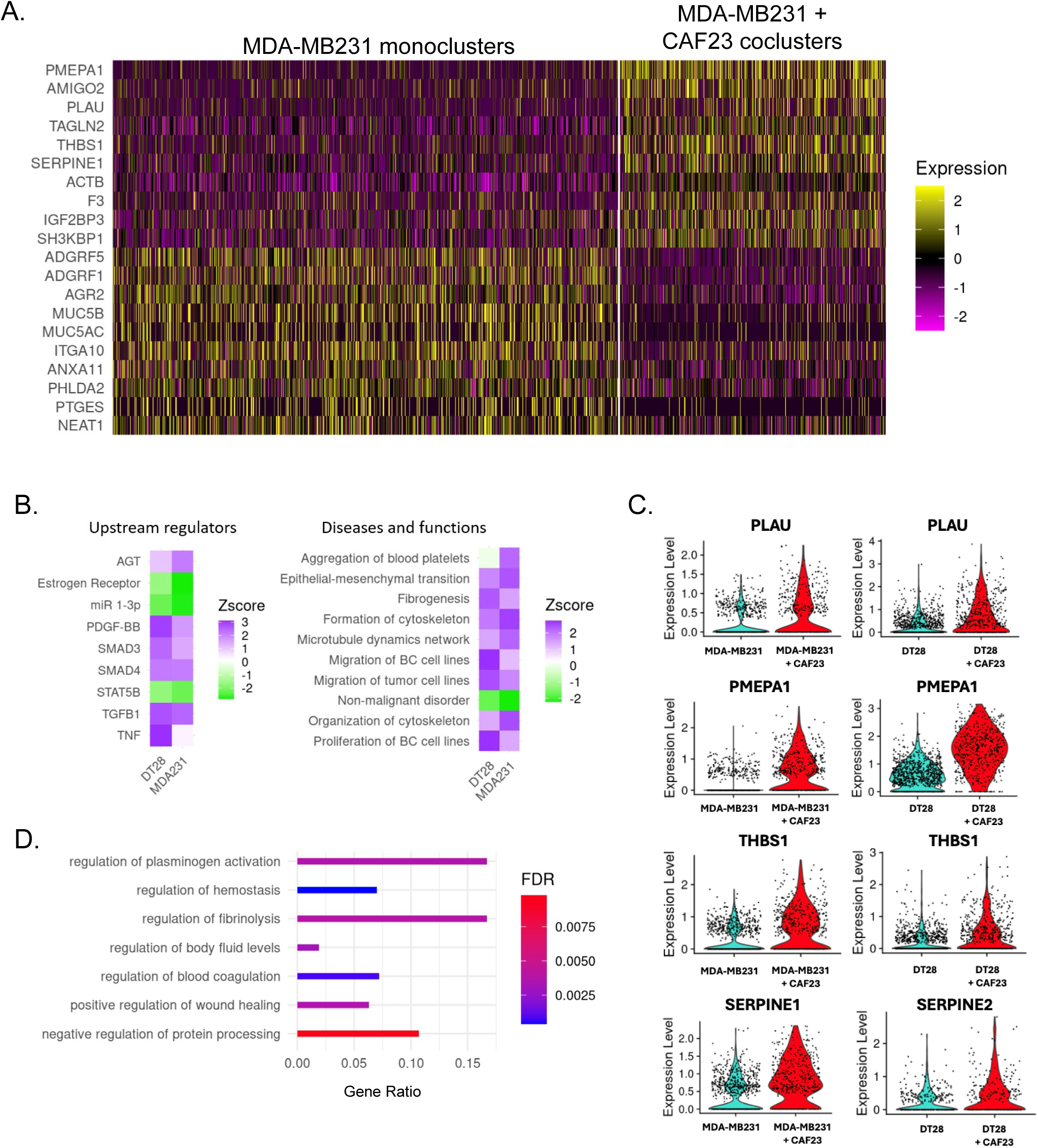
Single Cell RNA seq analysis of MDA-MB231 cells from MDA-MB231 mono-clusters vs CAF23/MDA-MB231 co-clusters. A. Heatmap showing the top 10 up and down DEG in breast cancer cells from MDA-MB231 mono-clusters vs breast cancer cells from CAF23/MDA-MB231 co-clusters. Plot shows only cells that were in the G1 phase of the cell cycle; each line represents an individual cell. Number of MDA-MB231 cells analyzed: 902 cells from MDA-MB231 mono-clusters and 474 cells from CAF23/MDA-MB231 co-clusters. B. Heatmap showing top upstream regulators and top diseases/functions identified by Ingenuity Pathway Analysis (IPA) of DEG lists of breast cancer cells from mono-clusters vs co-clusters for MDA-MB231 and DT28 cell lines. The Z score indicates the likely activation (positive) or inhibition (negative) of a regulator or function; a score of 2 is considered significant. C. Violin plots from the top 4 genes upregulated in both CAF23/MDA-MB231 co-clusters and CAF23/DT28 co-clusters. Each dot represents a cell. D. Gene Ontology analysis of a ranked list of DEG from MDA-MB231 mono-clusters vs CAF23/MDA-MB231 co-clusters. Gene ratio indicates the fraction of DEG genes over the number of genes in each gene set. FDR is the adjusted p value.

Using our list of DEG in co-clusters vs mono-clusters from both MDA-MB231 and DT28 samples we performed Ingenuity Pathway Analysis (IPA) with a cutoff of at least ± 0.25 Log2xFC and a Padj value of 0.05. Several upstream regulators were identified by IPA where at least one cell line had a Z-score of 2.0 or higher, including TGFβ1, PDGF-BB, AGT, and SMAD4 (Fig. 4B, left). Analysis of IPA Diseases and Functions identified gene sets such as epithelial-mesenchymal transition (EMT), formation of cytoskeleton, migration of breast cancer cell lines, and fibrogenesis (Fig. 4B, right). A small handful of top genes in these families were consistently identified, including uPA/PLAU, SERPINE1 or 2, THBS1, and PMEPA1 (Supplemental Fig. 5D). Violin plots (Fig. 4C) revealed that differences in these genes were readily apparent when viewed as individual cells. uPA/PLAU, SERPINE1/PAI1 or SERPINE2, THBS1, and S100A10 or S100A11 are all components of the plasminogen activation and fibrinolysis pathway[25]; Gene Ontology analysis (geneontology.org) confirmed this with top hits in these pathways and subfamilies of these pathways (Fig. 4D). Interestingly, while there were only 4-5 genes which were upregulated in BC cells from both DT28 co-clusters and MDA-MB231 co-clusters, similar upstream regulators and diseases/functions were identified for both types of TNBC cell lines (Fig. 4B). Both cell lines had estrogen receptor signaling downregulated. Both cell lines also had non-malignant disorder downregulated and EMT/migration upregulated, which is consistent with CAF cells promoting these hallmarks of cancer.

For the final analysis of our Single Cell RNA-seq data, we sought to understand if TNBC cells altered the transcriptional profile of CAF23 cells. For this, CAF23 mono-clusters were compared to each co-cluster type. HTOs again picked out the origin of each cell, and CAF markers were used to define whether cells were CAF23, MDA-MB231 or DT28s (Supplemental Fig. 6A). This time BC cells were removed so that CAFs could be compared to each other; after normalization and all other steps were performed, differential gene expression analysis generated a list of 300 DEGs that were differentially expressed in CAF23 cells when in CAF23/MDA-MB231 co-clusters and 113 genes in CAF23/DT28 co-clusters (with a Log2xFC of at least 0.5) (Supplemental Fig. 6B). Although genes induced by either MDA-MB231 or DT28 cells did not completely overlap, IPA analysis revealed similar canonical pathways were activated by either TNBC cell line, such as extracellular matrix organization, collagen biosynthesis, and integrin cell surface interactions (Supplemental Fig. 6C, left). Similar upstream regulators were also revealed, such as TGFβ, SMAD2, SMAD3, and AKT (Supplemental Fig. 6C, right). Overall, it appeared that clustering of CAFs and TNBC cells made the CAFs more CAF-like. This is perhaps not unexpected as CAFs cultured in isolation from breast cancer cells may have lost some (but certainly not all) extracellular matrix protein expression. When CAFs are put back together with cancer cells and exposed to signaling molecules secreted from BC cells that activate fibroblasts, such as TGFβ, this expression is restored.

Recombinant uPA or plasminogen urokinase, the central component of the plasminogen activation pathway, has been shown to bind to HUVEC cells[26]. It is encoded by the gene, PLAU. As uPA/PLAU was found upregulated in breast cancer cells from both MDA-MB231 and DT28 co-clusters (Fig. 4D), we focused on this protein as a potential mediator of BC-endothelial cell interaction. First, we sought to validate our Single Cell RNA seq. In immunofluorescence experiments, a uPA antibody stained positive in cells treated with PLAU siRNA (siPLAU) even though RTPCR confirmed that the knockdown was successful (Fig. 5B). Western blots showed that a 48 kDa band (the size of uPA) disappears in siPLAU cells (Fig. 5B inset); a 66 kDa band, the size of tissue specific plasminogen activator (tPA), does not change (Supplemental Fig. 7A, B). Because of the non-specificity of the uPA antibody, we were unable to perform immunofluorescence experiments with BC/CAF co-clusters. Instead, we tested whether CAF conditioned media (CM) could induce uPA/PLAU. 2D-grown MDA231, DT28, and SUM159 cells were starved for 24 hrs in 1% DCC and then treated for 24 hrs in either CAF CM +1% DCC or IMEM + 1% DCC. DCC (charcoal-stripped media) was used as a source of serum in order to eliminate effects of steroid hormones. We observed that CAF CM was able to induce uPA/PLAU (48 kDa band, Fig. 5A) in MDA-MB231 cells. Although we did not observe induction of uPA/PLAU in DT28 or SUM159 cells, we tested whether knockdown by siPLAU could affect endothelial binding in all three cell types. In MDA-MB231 cells treated with siPLAU for 3 days before the endothelial binding assay, we observed a significant decrease in endothelial binding that ranged from 40-60% (Fig. 5C). The effect in DT28 and SUM159 cells was less intense but still statistically significant. When the assay was performed as a time course over 40 minutes, a 50% drop in endothelial binding with siPLAU was consistently observed at every time point (Fig. 5D). PLAU siRNA did not affect the ability of MDA-MB231 to form co-clusters with CAF23 cells (Supplemental Fig. 7C). uPA is a serine protease that converts plasminogen to its active form, plasmin. The plasminogen activator system activates downstream metalloproteases such as MMP2 and MMP9, which promote degradation of basement membrane and ECM proteins during cancer cell invasion and migration[25]. We next sought to determine if uPA’s serine protease activity was necessary for binding to endothelial cells by utilizing a uPA specific inhibitor, UK122[27]. MDA-MB231 cells were pre-treated with UK122 for 24 hrs to allow time for the absence of uPA activity to influence ECM organization. In the endothelial binding assay, cells pre-treated with 100 µM UK122 bound to endothelial cells with significantly less efficiency (Fig. 5E); however, binding was not as severely affected as it was with PLAU siRNA knockdown experiments.

**Figure 5.**
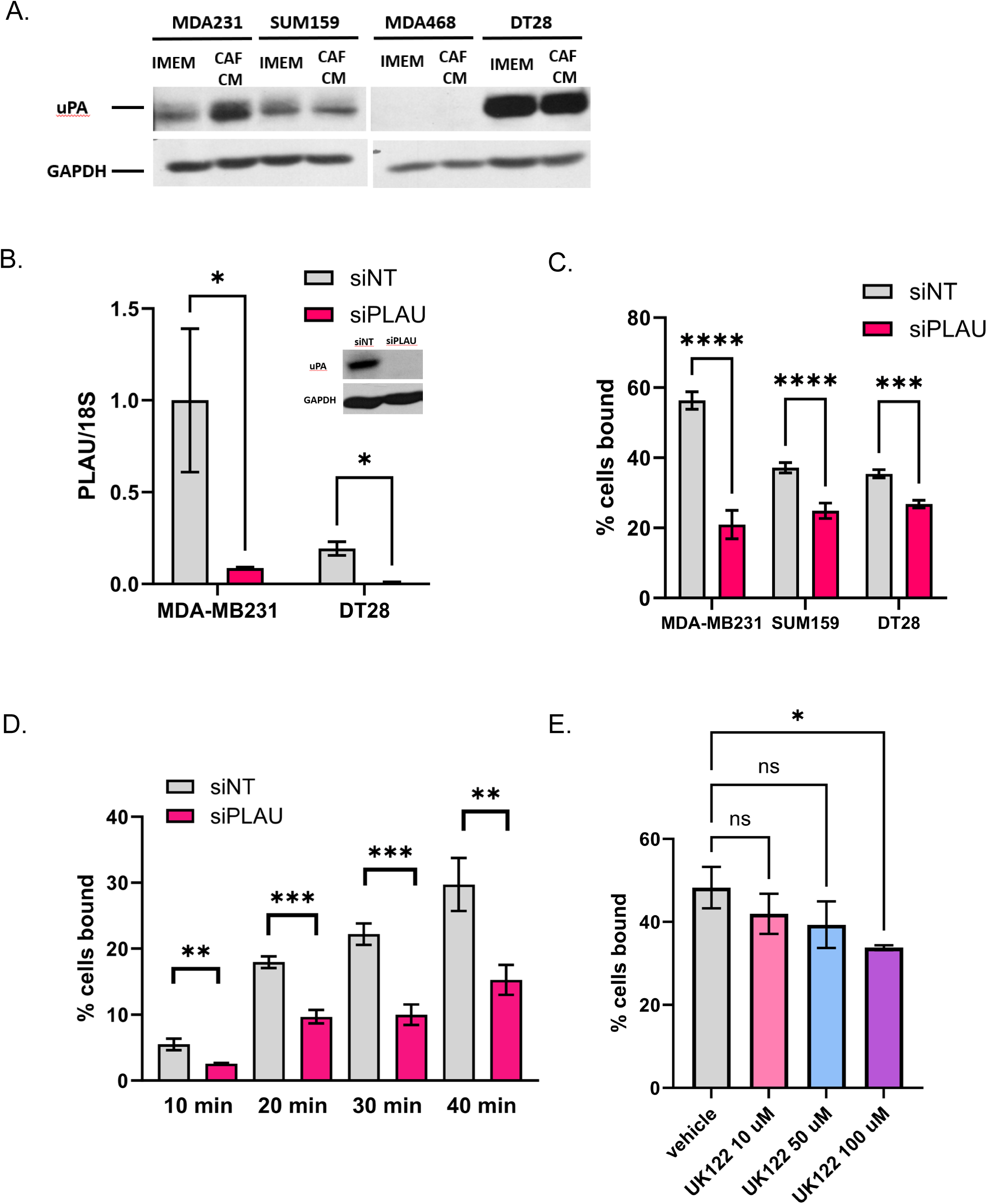
Reduction of uPA/PLAU inhibits BC-endothelial cell binding. A. Western blot showing that CAF23 conditioned media increases uPA protein in MDA-MB231 cells. B. qPCR of PLAU in MDA231 or DT28 cells treated for 3 days with non-targeting siRNAs (siNT) or PLAU siRNAs (siPLAU). PLAU was normalized to the 18S ribosomal RNA. Inset: Western blot of siNT or siPLAU MDA-MB231 cells. C. Endothelial binding assay of MDA-MB231, DT28 and SUM159 cells treated for 3 days with siNT or siPLAU. D. Time course of endothelial binding assay using MDA-MB231 cells treated with siNT or siPLAU showing % of cells bound for the indicated periods. E. Endothelial binding assay using MDA-MB231 cells pre-treated with vehicle or uPA inhibitor, UK122, for 24 hrs at increasing concentrations. * p ≥ 0.05, ** p ≥ 0.005, *** p ≥ 0.0005

uPA/PLAU is regulated by multiple secreted factors, including TGFβ[28], TNFα, VEGF and FGF2. As TGFβ was identified as an upstream regulator in CAF/BC co-clusters (Fig. 4B), we next sought to determine the effects of TGFβ on endothelial binding. RTPCR confirmed that BC cells exposed to TGFβ increases PLAU transcripts (Supplemental Fig. 8A). We first tested the immediate effects of TGFβ signaling on endothelial binding by treating cells with TGFβ for 1 hr before the endothelial binding assay; we saw a small but insignificant enhancement of binding both with siNT and siPLAU treated cells (Supplemental Fig. 8B). Additional experiments with a longer, 24 hr TGFβ treatment did not show any enhancement in endothelial binding (Supplemental Fig. 8C). One explanation for this is that TGFβ regulates a number of genes, including SERPINE1 (PAI1)[28], a negative regulator of uPA. SERPINE1 was also identified as an upregulated gene in our Single Cell RNA seq analysis (Fig. 4D). CAFs secrete of a number of different factors which could influence endothelial binding. Therefore, direct application of TGFβ alone may not replicate the effects of CAFs on breast cancer-endothelial cell interactions.

We next determined if PLAU siRNA on MDA-MB231 cells could affect extravasation in the ex vivo assay. First, we confirmed by RTPCR that siPLAU remained effective up to 48 hrs after it is removed (Fig. 6B), thus giving us a sufficient time window to evaluate the effects of siPLAU in an ex vivo assay. We therefore made 24-hr siNT and siPLAU MDA-MB231 mono-clusters (cells were grown cells with siRNAs for 2 days under adherent conditions and for 1 extra day under clustering conditions for 3 days total siRNA exposure) and performed the ex vivo extravasation assay as outlined (Fig. 6A). We did not observe any difference in early times of extravasation (8 and 24 hrs), when MDA-MB231 mono-cluster extravasation is still typically low (Fig. 2C). However, at 48 hrs, when 45% of siNT MDA-MB231 mono-clusters had extravasated, we saw that only 25% of siPLAU MDA-MB231 had extravasated (Fig. 6C). Thus, we conclude that uPA/PLAU enhances BC cell extravasation.

**Figure 6.**
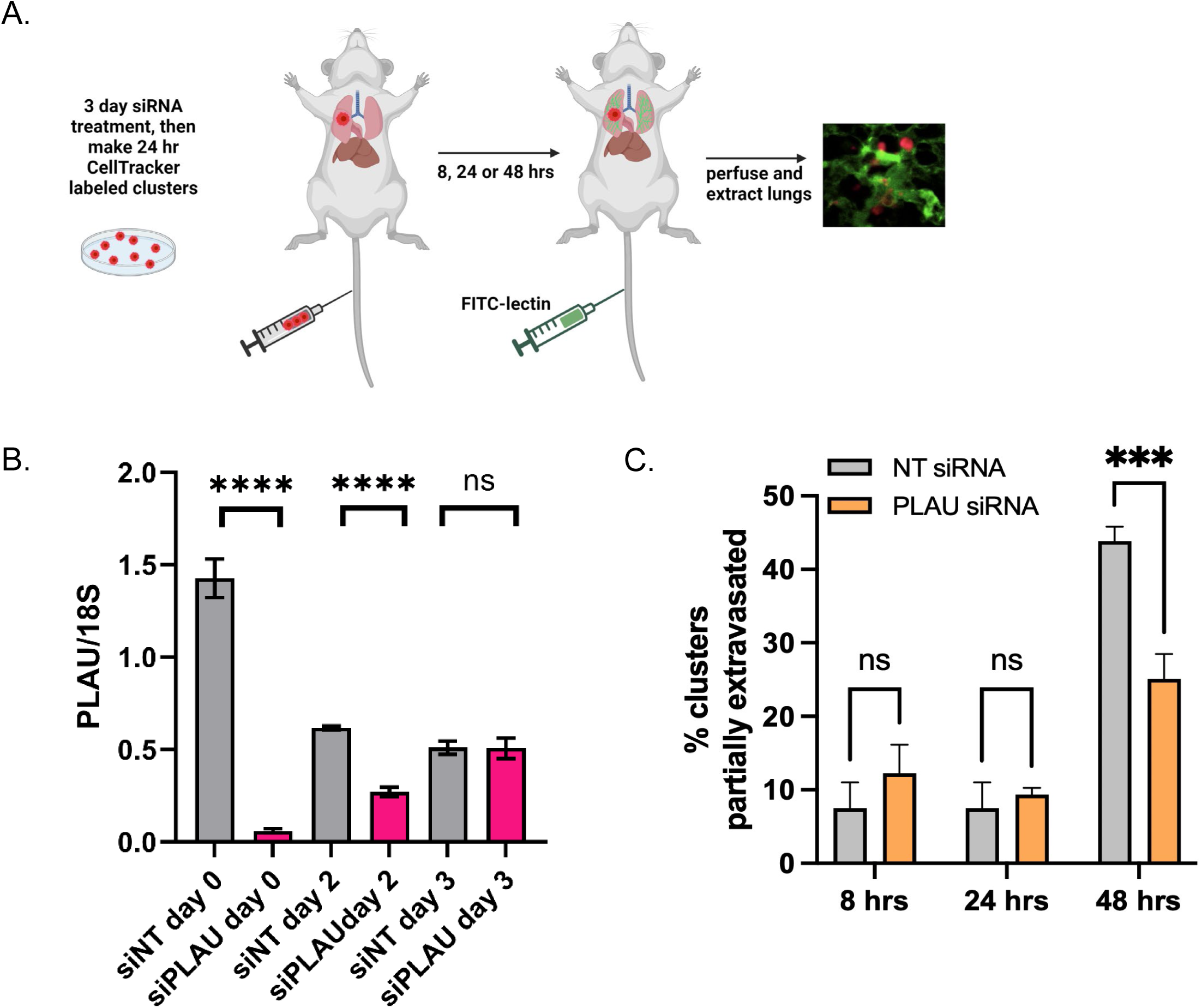
Reduction of uPA/PLAU in MDA-MB231 cells inhibits extravasation in the ex-vivo lung assay. A. Schematic of ex vivo assay. MDA-MB231 cells were treated with siNT or siPLAU for 2 days, then cells were stained with CellTracker Red and made into 24 hr mono-clusters in the presence of siNT or siPLAU (3 days total siRNA exposure). Clusters were collected and injected by tail vein into FVB mice. 30 minutes prior to lung perfusion, mice were injected by tail vein with FITC-lectin to stain the vasculature. Lung perfusion and fixation was performed at 8 hrs, 24 hrs, and 48 hrs. Pieces of lung were cut and imaged with confocal microscopy to visualize and quantify BC clusters inside or outside of the vasculature. B. qPCR experiment showing persistence of siRNA effect. MDA-MB231 cells were treated with siRNAs for 3 days as before. Cells were then washed of siRNA, cultured in regular media, and qPCR was performed at the given times after removal of siRNA. 18S was used as a control for the qPCR. C. Results from ex vivo assay described in A using 2 mice per time point, per treatment. 20-40 clusters were counted for each animal and scored as being completely intravascular or partially extravascular.

## Discussion

Cancer-associated fibroblasts, as the most abundant non-cancerous cell type in the tumor microenvironment, have been implicated in nearly all of the described hallmarks of cancer, most often (though not exclusively) playing a pro-tumorigenic role. Numerous studies have investigated their influence on cancer cell invasion, migration, conversion to an EMT status, and stemness[4–6], all processes that take place when cancer cells are in a stable environment, growing and residing within tissues. Conversely, we have focused on how CAFs influence breast cancer cells when they are present in the volatile environment of the blood circulation as circulating CAFS (cCAFS) or as cCAF/CTC co-clusters. Here we have presented another pro-tumorigenic function of CAFs, that of cancer cell extravasation from the circulation, a necessary step in the metastatic cascade.

We propose three ways that CAFs might aid breast cancer cell extravasation. First, due to direct contact, CAFs may lead breast cancer cells across the endothelial cell layer (Fig. 7, top). We observed that CAFs bind better to endothelial cells than breast cancer cells, and therefore they may have a more profound influence on endothelial cell interactions than breast cancer cells alone. A potential candidate to mediate CAF-endothelial cell interactions is CD44; the sialofucosylated glyco form of CD44 is known as HCELL and interacts with endothelial E-selectin during the tethering step of the rolling model of extravasation[18]. CD44 is more highly expressed in CAFs than in MDA-MB231 cells. We did not observe an effect of CD44 siRNA in our endothelial binding assay for either MDA-MB231 or CAF23 cells; however, this could be due to the preparation of our HUVEC cell lawn as we did not stimulate expression of E-selectin in our assay. Regardless, CD44 is important for mediating interactions between CAFs and BC cells, as we have already shown[8, 9]. Hyaluronin, or HA, is a CD44 ligand that we also tested. Depletion of HA significantly decreased CAF-endothelial binding.

**Figure 7.**
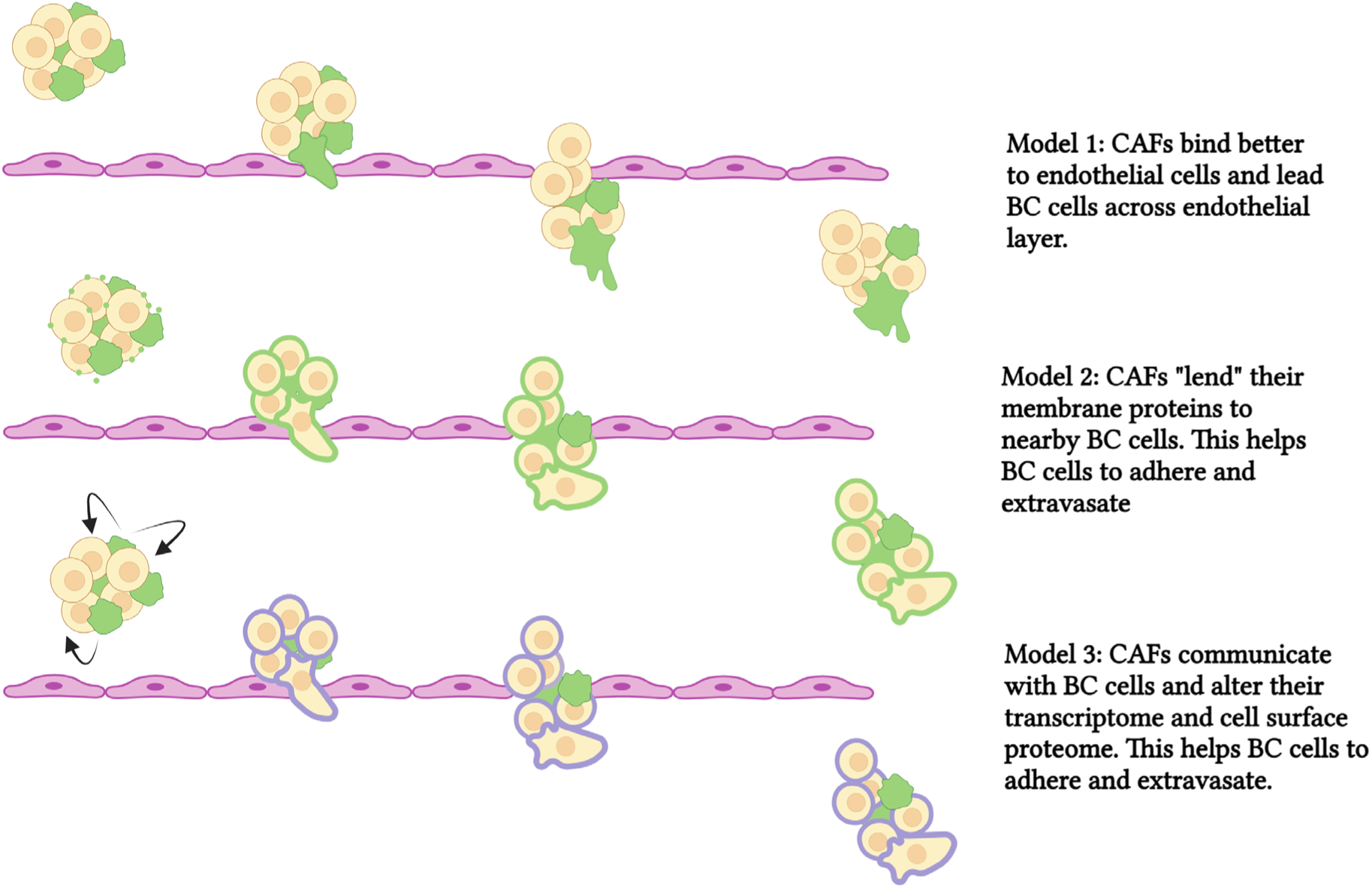
Schematic of models showing CAF-aided BC extravasation. In the top panel, CAFs bind to vascular endothelium better than breast cancer cells. CAFs then lead BC cells across the endothelium. In the middle panel, CAFs lend ECM proteins to BC cells (by secreted extracellular vesicles) and this increases endothelial binding and promotes extravasation. In the bottom panel, CAFs communicate with BC cells and alter the BC transcriptome. This increases proteins, such as uPA, that promote extravasation. CAFs are depicted in green, breast cancer cells are depicted in beige/yellow. ECM proteins on CAFs are in green. Proteins induced by CAFs and expressed in breast cancer cells in purple. *** p ≥ 0.0005. **** p ≥ 0.00005

CTCs interact with several different types of other cells within the circulation that may aid CTC extravasation. Soon after intravasation, CTCs attract and activate platelets, which surround them to form microthrombi. Platelets perform many functions that help CTCs to survive in the vasculature, such as protecting CTCs from shear forces and natural killer cells[14, 29]. Platelets express many types of integrins, which as discussed in the introduction, form firm adhesions with endothelial cells. Thus, platelets may act as linker cells between endothelial cells and CTCs to mediate TEM. Additionally, platelets express a complex mixture of growth factors, chemokines, and cytokines (they are the major source of TGFβ in the circulation) and attract other cell types such as neutrophils, monocytes, or leukocytes which may also act as linker cells. Macrophages also play a prominent role in CTC extravasation[30–32]. Through lung intravital imaging in mouse models, Genna et al. demonstrated the presence of physical interactions across the vascular barrier between CTCs and macrophages via thin membranous extensions[33]. The precedent of platelets and macrophages acting as linker cells suggests that CAFs too, may act as linker cells. Real-time intravital imaging would be informative to definitively record the progression of CAF-mediated extravasation and their interactions with breast cancer cells, endothelial cells, and other cell types that are recruited to sites of extravasation.

A second way that CAFs might influence cancer cell extravasation is through sharing of proteins, DNA, lipids, and micro-RNAs via secretion of extracellular vesicles (EVs) and subsequent uptake by cancer cells (Fig. 7, middle). Extracellular vesicles consist of microsomes (30-100 nm, arising from endosomal compartments within the cell), and microvesicles (100 nm to 1 µm, which bud off from cellular membrane). Intercellular CAF-cancer cell communication through EVs has been documented to have effects on proliferation, migration, invasion, and even chemoresistance[34]. Santi et al. used an isotope-labeling/mass spec method to identify CAF-derived proteins that were transferred to endothelial cells when CAFs and HUVECs were co-cultured, and one prominent protein they identified from this study was CD44[23]. We have previously observed that some MDA-MB231 cells treated with CD44 siRNA stained positively for CD44 when co-cultured with CAF23 cells, suggesting that CD44 was imparted onto the breast cancer cells from CAFs (data not shown). As mentioned above, we have not ruled out CD44 as having an influence on MDA-MB231-endothelial cell interactions. Other proteins that could affect endothelial binding through EV secretion from CAFs include hyaluronin, integrins, or other membrane proteins. It should be noted that the CAF23 conditioned media used in Figure 3E was filtered through a 0.2 micron membrane that would have excluded EVs and microsomes, leading us to conclude that increased endothelial binding in our studies was due to secreted, soluble factors. Hyaluronin is both membrane-associated and soluble.

Finally, as we show with our Single Cell RNA seq data, CAFs communicate with breast cancer cells when they are in close contact. CAFs are hypersecretory[4–6], and we have shown previously that our CAF23 cells secrete numerous factors including TGFβ, IL-8, IL-6, CXCL12/SDF1, and FGF2 ([9] and data not shown) that may be received as signals to alter the transcriptome, and ultimately the membrane proteome or extracellular matrix (ECM), of BC cells to increase their interaction with mammary endothelial cells. Thus, CAFs may indirectly increase breast cancer cell-endothelial cell interactions to facilitate extravasation (Fig. 7, bottom). This could be significant because at the time when cancer cells are ready to extravasate, CAFs and BC cells may have become dissociated as the cell clusters are squeezed into small vessels and capillaries, as we observed in our ex vivo assays.

Our Single Cell RNA-seq data revealed that CAFs increase a number of genes (PLAU, SERPINE1/PAI1, and THBS1) that are upregulated by TGFβ. We chose to focus on PLAU and were able to implicate uPA/PLAU in breast cancer extravasation as seen by the in vitro endothelial binding assay and the ex vivo lung extravasation assay. The PLAU gene encodes the urokinase-type plasminogen activator (uPA), a serine protease which becomes active upon binding to its receptor (uPAR), a GPI-anchored membrane protein, and catalyzes the conversion of plasminogen to plasmin. Plasminogen and plasmin are components of the fibrinolytic system that dissolves blood clots. Plasmin itself is a protease that activates growth factors and other proteases such as MMP-2 and MMP-9 within the ECM. Hence uPA sets off a protease cascade important for ECM reorganization and therefore is a major player in cancer cell migration, invasion, EMT and stemness[25, 35, 36]. uPA is regulated by the plasminogen activator inhibitor, SERPINE1(PAI1). uPA, uPAR, and PAI1 have been correlated with poor progression in many cancers including breast, ovarian, pancreatic, colorectal, and lung cancer[37, 38]. We found that uPA was upregulated by CAFs in CAF23/MDA-MB231 and CAF23/DT28 co-clusters while PAI1 was also upregulated in CAF23/MDA-MB231 co-clusters. SERPINE2 is a lesser known uPA inhibitor that was upregulated in CAF23/DT28 co-clusters. uPA and its inhibitor are both implicated in pro-tumorigenic processes; while seemingly contradictory, this paradox is widely known[39]. uPA and PAI1 have been used for decades as biomarkers in diagnostic assays for breast cancer patients, and the ratio of uPA/PAI1 appears to be more prognostic than either protein by itself[40].

uPA, and its receptor UPAR, have diverse functions that could separately or cumulatively affect extravasation. Barnathan et al. showed that recombinant single chain uPA (pro-uPA) or its amino terminal fragment, ATF (aa 1-143), containing the UPAR-binding domain, bind directly to HUVEC cells[26]. uPA ATF lacks the enzyme’s enzymatic activity domain and has lower fibrinogen turnover, demonstrating that uPA proteolytic activity is not necessary for all of the functions of uPA. In our endothelial binding assays, we saw a reduction in binding with the uPA inhibitor, UK122, but it was not as effective as siPLAU. This suggests that uPA protease activity has some effect on binding of breast cancer cells to endothelial cells, but it is not exclusively required. uPA enhances binding of its receptor, UPAR, to integrin receptors, vitronectin (an ECM protein), formyl peptide receptors, and growth factor receptors (such as EGFR or PDGFR), with resulting effects on cell adhesion, migration, invasion, and proliferation[25, 36, 37]. Our assays were conducted after several days of siRNA treatment (or after 24 hr UK122 treatment) which allowed time for changes in the composition of the BC ECM to take place but also allowed time for inhibition of uPA/UPAR downstream pathways to be affected. A close examination of the CAF23/BC ECM proteome would provide an interesting perspective on how CAFs may affect BC cell-endothelial cell interactions.

The consequences of CAFs in the circulation are multi-fold. While we have focused the presentation of our Single Cell RNA seq data on the candidates that may be involved in extravasation, we also found that DEG in breast cancer cells regulated by CAFs (PMEPA1, uPA, PAI1, THBS1, F3) are components of EMT, migration, and proliferation pathways. The discovery of the plasminogen urokinase system being upregulated by CAFs is especially significant as uPA/UPAR activates downstream signaling pathways such as integrin receptor and growth factor receptor signaling[36]. Thus, while CAF-mediated extravasation of breast cancer cells would result in increased survival of breast cancer cells, CAF/BC cell interactions also aid the establishment and proliferation of metastatic BC cells once they extravasate. The importance of circulating CAF/BC cell clusters cannot be understated, and investigation of these cellular interactions in the context of the circulation should be studied further. A more complete understanding of the mechanisms of these processes will reveal novel ways to target the CAF/CTC compartment to interrupt breast cancer dissemination and prevent new metastases.

Single Cell RNA seq data will be deposited into the Gene Expression Omnibus database once the paper has been submitted for publication in a peer-reviewed journal.

## Acknowledgments

This work was supported by the Breast Cancer Research Foundation (Project 00086668 to CAL), National Institutes of Health (NIH) (R01 CA229697 to CAL), Department of Defense (DOD W81XWH-19-1-0699 to DEA and CAL), and the Tickle Family Land Grant Endowed Chair in Breast Cancer Research (to CAL). Imaging and RNA sequencing was supported by resources and staff at the University of Minnesota Genomics Core (UMGC) and the University Imaging Centers (UIC).

