## Supplemental Figures for "Modeling the role of urokinase plasminogen activator, uPA, and circulating Cancer-Associated Fibroblasts (cCAFS) in breast cancer cell extravasation"

Supplemental Figure 1: **Perturbing CD44/hyaluronin axis in the endothelial binding assay**. A. Immunofluorescence experiments with anti-CD44 showing siRNA knockdown of CD44 or a non-targeting siRNA control in CAF23 and MDA-MB231 cells. B. Endothelial binding assay with the cells in A. C. Immunofluoresence experiments with HA-binding protein/anti-streptavidin in CAF23 and MDA-MB231 cells that were treated overnight with 4MU and for 1 hr with hyaluronidase to deplete HA protein. D. Endothelial binding assays with the cells in C. ** $p \geq 0.005$, **** $p \geq 0.00005$

Supplemental Figure 2. **Single Cell RNA seq analysis of CAF23 single cells grown in 2D**. A. FeaturePlots showing the expression of CAF markers commonly described in the literature on CAF23 single cells grown under adherent conditions. Total number of cells analyzed = 3884.

Supplemental Figure 3. **Single Cell RNA seq analysis of CAF23 single cells compared to MDA-MB231 single cells.** A. Dim plot showing CAF23 and MDA-MB231 cells based on their HTO classification during demultiplexing. Number of cells analyzed: CAF23 = 3766, MDA-MB231 = 1491. B. Feature plots showing expression of CAF markers in the cells that correlate with CAF23 cells in A. C. Heat map showing top 20 up and down DEG in CAF23 vs MDA-MB231 cells grown under 2D adherent conditions. Only features that had a minimum percent difference of 0.65 were calculated for this plot to select markers that had extremely low levels in MDA-MB231 cells. Each line represents a cell. D. Line plot showing the top hits from the Cellular Component gene sets in Gene Ontology analysis using a DEG list from CAF23 cells compared to MDA-MB231 cells. This analysis indicates that CAF23 cells are likely matrix CAFs (mCAFs). Count is the number of genes in the DEG list for any particular GO gene set. Gene ratio is the fraction of DEG genes in the list from the total number of genes in the GO gene set. FDR is the adjusted P value.

Supplemental Figure 4: **Single Cell RNA seq analysis of MDA-MB231 cells from MDA-MB231 mono-clusters vs CAF23/MDA-MB231 co-clusters.** A. Dim plot showing MDA-MB231 mono-clusters and CAF23/MDA-MB231 co-clusters based on their HTO classification during demultiplexing (left) and feature plots showing expression of CAF markers (right). CAF23 cells are highlighted. These cells are removed in downstream analysis (shown in Figure 4). B. Dot plot showing the top DEG from breast cancer cells from MDA-MB231 mono-clusters vs CAF23/MDA-MB231 co-clusters, the average expression value of all cells, and the percentage of cells that are expressing. For this plot, DEG that were expressed in 100% of cells were excluded.

Supplemental Figure 5: **Single Cell RNA seq analysis of DT28 cells from DT28 mono-clusters vs CAF23/DT28 co-clusters.** A. Dim plot showing DT28 mono-clusters and CAF23/DT28 co-clusters based on their HTO classification during demultiplexing (left) and feature plots showing expression of CAF markers (right). CAF23 cells are highlighted. These cells are removed in downstream analysis. B. Heatmap showing the top 20 up and down DEG in breast cancer cells from DT28 mono-clusters vs CAF23/DT28 co-clusters. Plot shows only cells that were in the G1 phase of the cell cycle. Number of DT28 cells analyzed: DT28 mono-clusters = 985, CAF23/DT28 co-clusters = 585. Each line represents a cell. C. Dot plot showing the top DEG in breast cancer cells from DT28 mono-clusters vs CAF23/DT28 co-clusters, the average expression value of all cells, and the percentage of cells that are expressing. For this plot, DEG that were expressed in 100% of cells were excluded. D. Heat maps showing comparison of MDA-MB231 and DT28 datasets for genes in the top hits from IPA. Expression log ratio is the Log2xFC value from the DEG analysis and indicates the change from breast cancers cells from mono-clusters vs co-clusters. Only the top up or down genes are shown.

Supplemental Figure 6: **Single Cell RNA seq analysis of CAF23 cells from CAF23 mono-clusters vs CAF23/MDA-MB-231 co-clusters or CAF23/DT28 co-clusters.** A. Dim plot showing cells from CAF23 mono-clusters, CAF23/MDA-MB231 co-clusters, and CAF23/DT28 co-clusters based on their HTO classification during demultiplexing (left) and feature plots showing expression of CAF markers (right). CAF23, DT28, and MDA-MB231 cells are highlighted. The BC cells are removed in downstream analysis. Number of CAF23 cells analyzed: CAF23 mono-clusters = 589, CAF23/DT28 co-clusters = 483, CAF23/MDA-MB231 = 314. B. Heatmap showing the top 30 genes that are upregulated in CAF23 cells from CAF23/BC-co-clusters compared to CAF23 mono-clusters. Each line represents a cell. C. Heatmap showing top canonical pathways and upstream regulators identified by IPA of DEG lists comparing CAF23 cells from mono-clusters vs CAF23 cells from CAF23/MDA-MB231 co-clusters or CAF23/DT28 co-clusters. The Z score indicates the likely activation (positive) or inhibition (negative) of a pathway or regulator; a score of 2 is considered significant.

Supplemental Figure 7: **Additional PLAU analysis**. A. Full western blot of a repeat experiment of that shown in Figure 5A. B. Additional western blot showing the absence of uPA in MDA-MB231 cells that were treated with siPLAU. For A and B, additional band shown at 61 kDa might be tPA. C. Clustering assay of siPLAU or siNT treated MDA-MB231 cells with CAF23 cells.

Supplemental Figure 8: **Effect of TGFβ on uPA/PLAU mediated endothelial binding**. A. qPCR analysis showing induction of PLAU in MDA-MB231 cells that were treated with 1-10 μg/ml recombinant TGFβ for 1 hr. B. Endothelial binding assay of siPLAU or siNT MDA-MB231 cells that were treated with 10 μg/ml TGFβ for 1 hr prior to the binding assay. C. Same as B, but cells were treated with 10 μg/ml TGFβ for 24 hrs (while still in the presence of siRNAs) prior to the endothelial binding assay. 18S was used as a control for the qPCR. **** $p \geq 0.00005$

### Supplemental Table 1

Supplemental Table 1: Hashtag oligos used in single cell RNAseq experiments

| Hashtag Oligo name (Biolegend Cat #) | Labeled cell population | Sequence |
|---|---|---|
| Experiment 1 | | |
| HTO6 (A0256) | MDA-MB231 single cells | GGTTGCCAGATGTCA |
| HTO7 (A0257) | CAF23 single cells | TGTCTTTCCTGCCAG |
| HTO8 (A0258) | MDA-MB231 monoclusters | CTCCTCTGCAATTAC |
| HTO10* (A0260) | CAF23/MDA-MB231 co-clusters | ATTGACCCGCGTTAG |
| Experiment 2 | | |
| HTO10* (A0260) | DT28 single cells | ATTGACCCGCGTTAG |
| HTO1 (A0251) | DT28 monoclusters | GTCAACTCTTTAGCG |
| HTO2 (A0252) | CAF23/DT28 co-clusters | TGATGGCCTATTGGG |
| HTO3 (A0253) | CAF23 monoclusters | TTCCGCCTCTCTTTG |

* HTO10 was used in both Experiment 1 and Experiment 2. Since these were contained in different HTO libraries, we were able to separate them with Seurat during data analysis.

Supplemental Figure 1

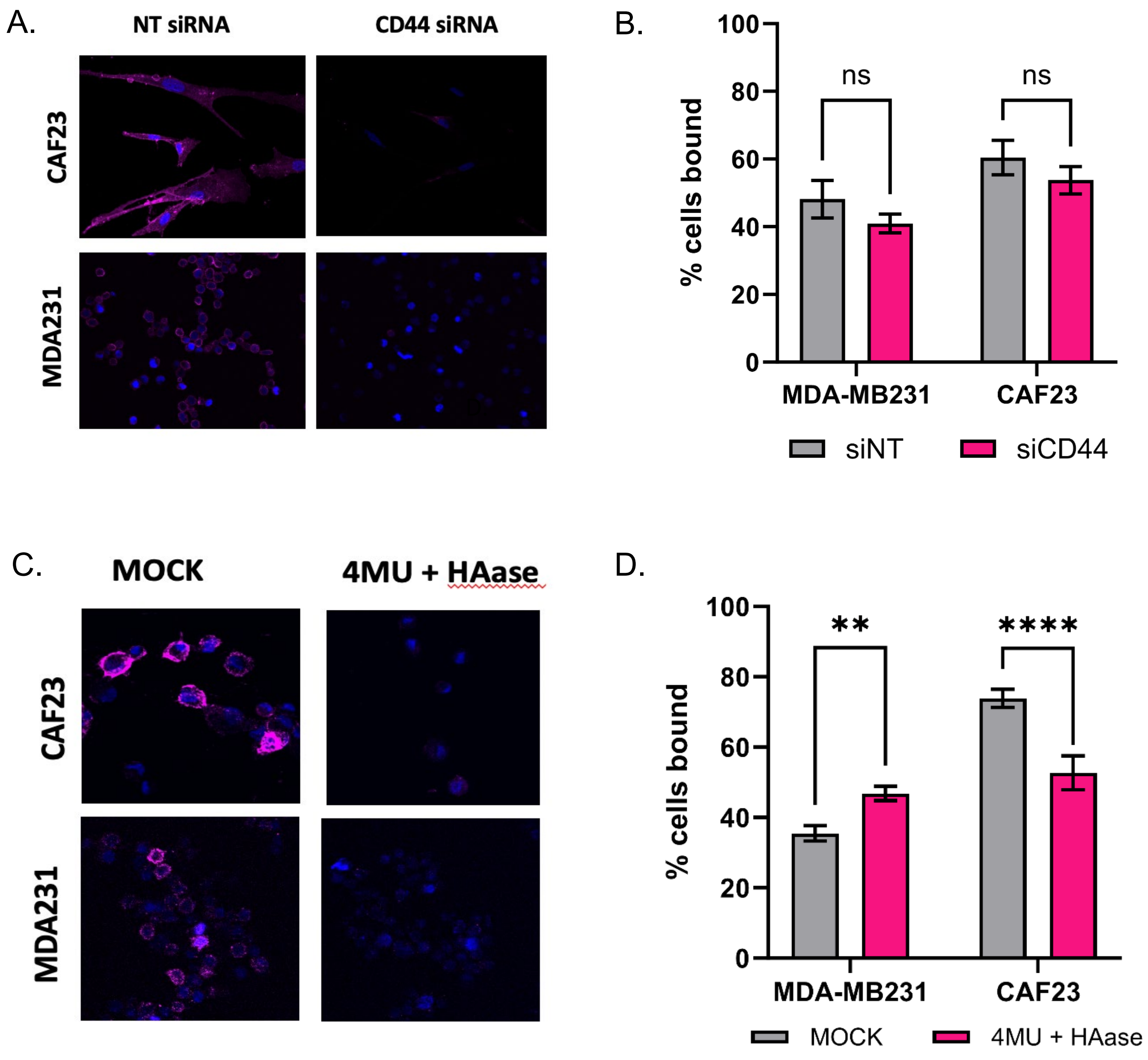

Supplemental Figure 2

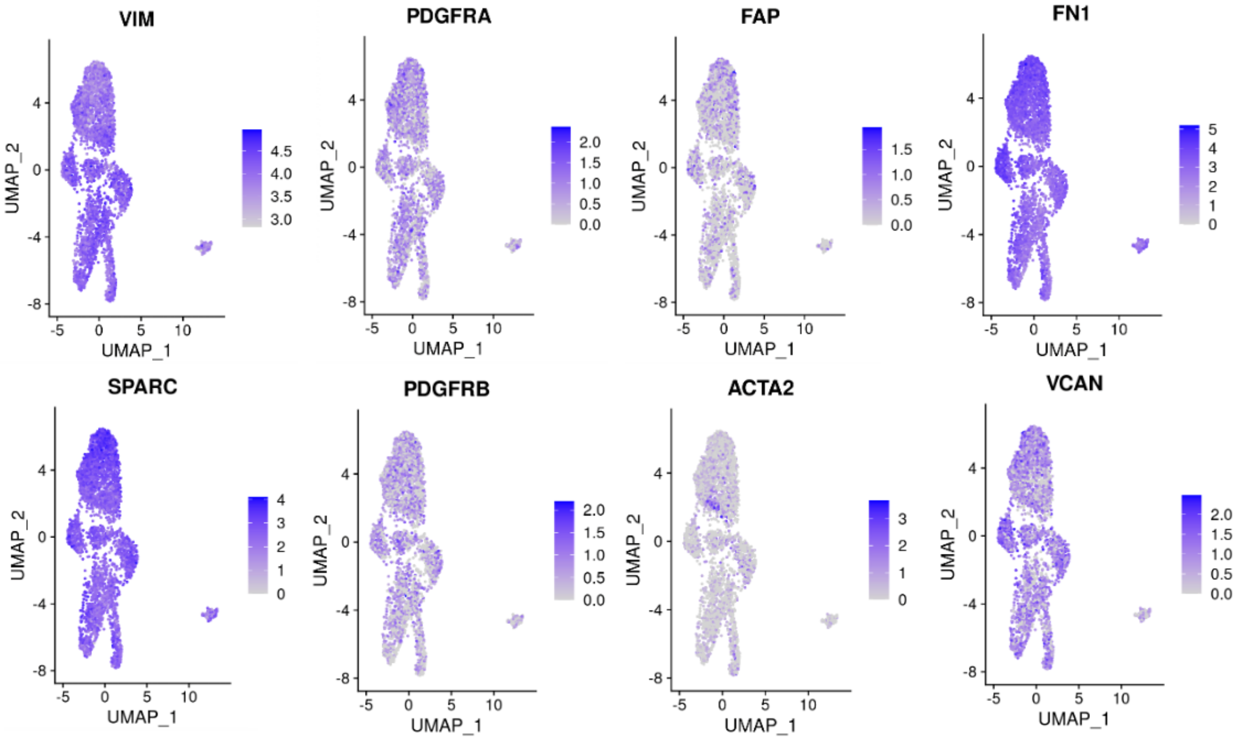

Supplemental Figure 3

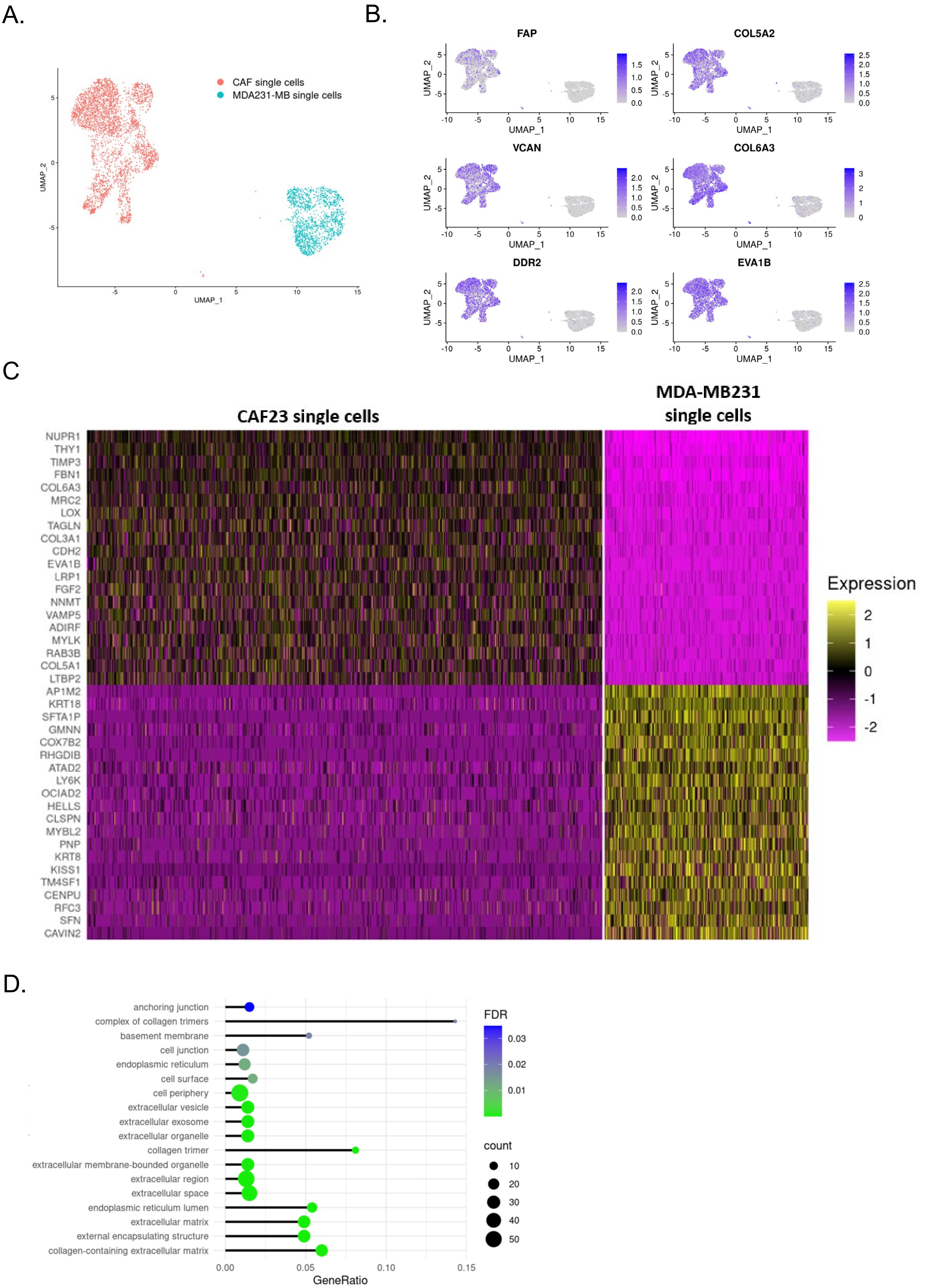

Supplemental Figure 4

A.

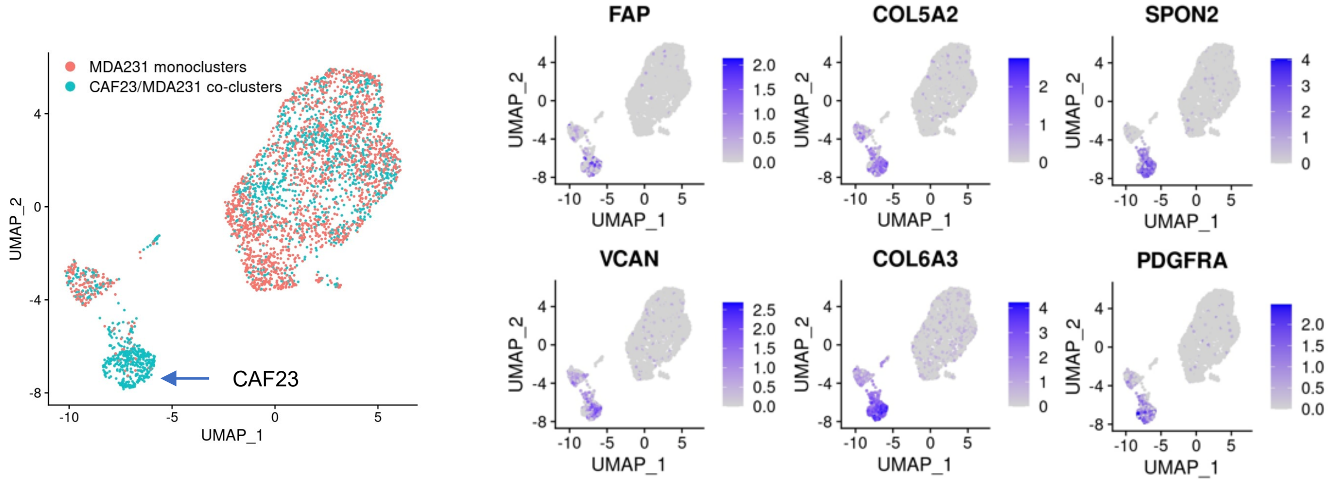


B.

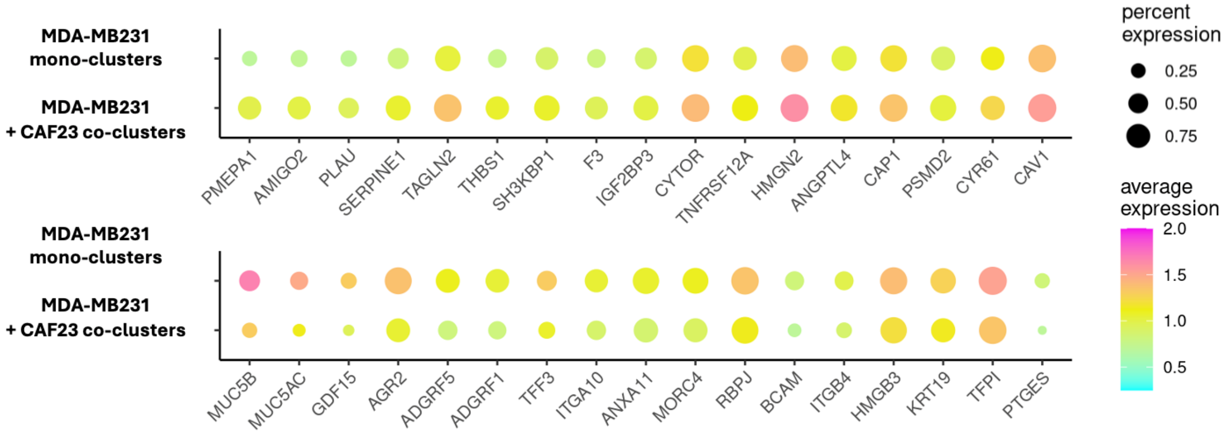

### Supplemental Figure 5

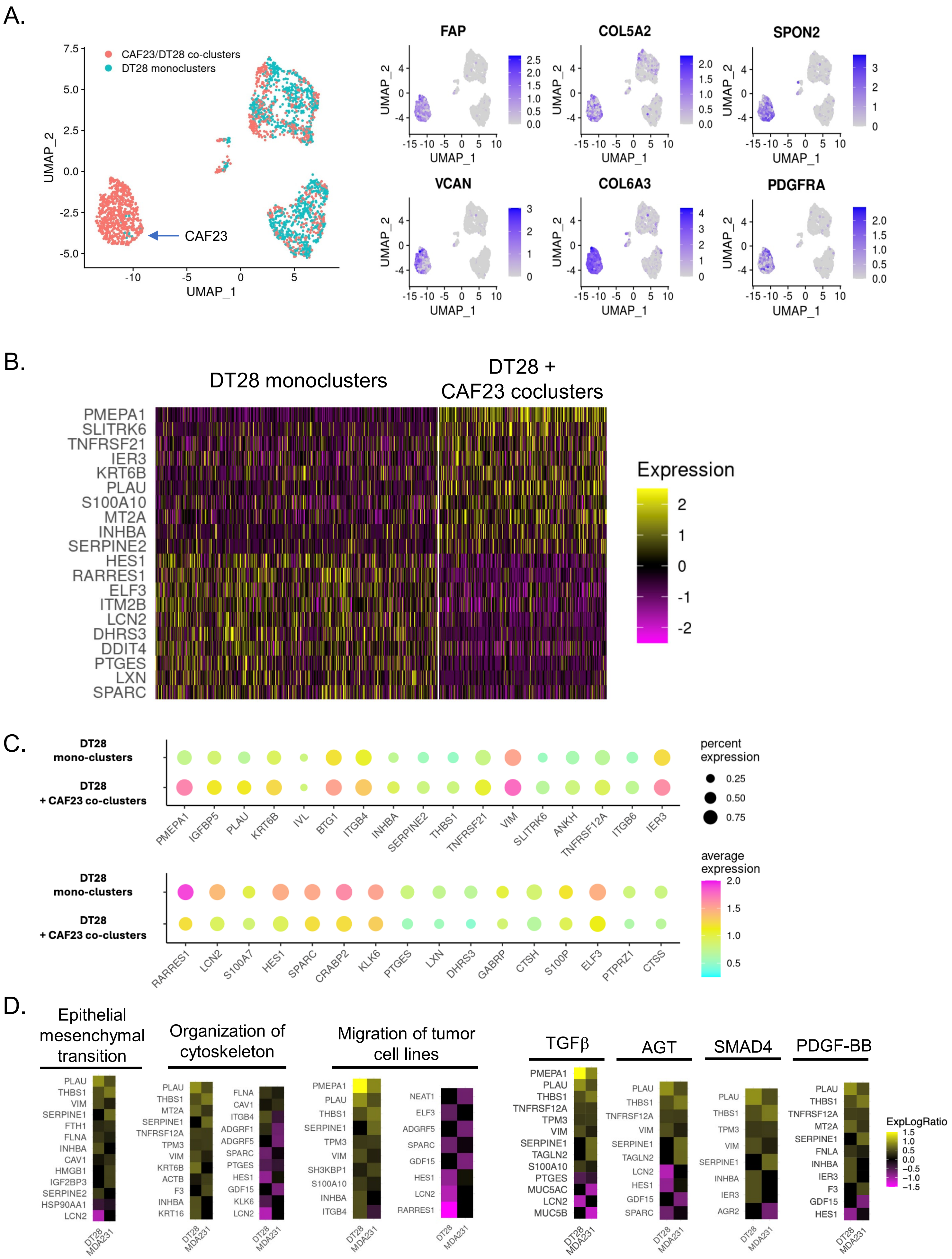

Supplemental Figure 6

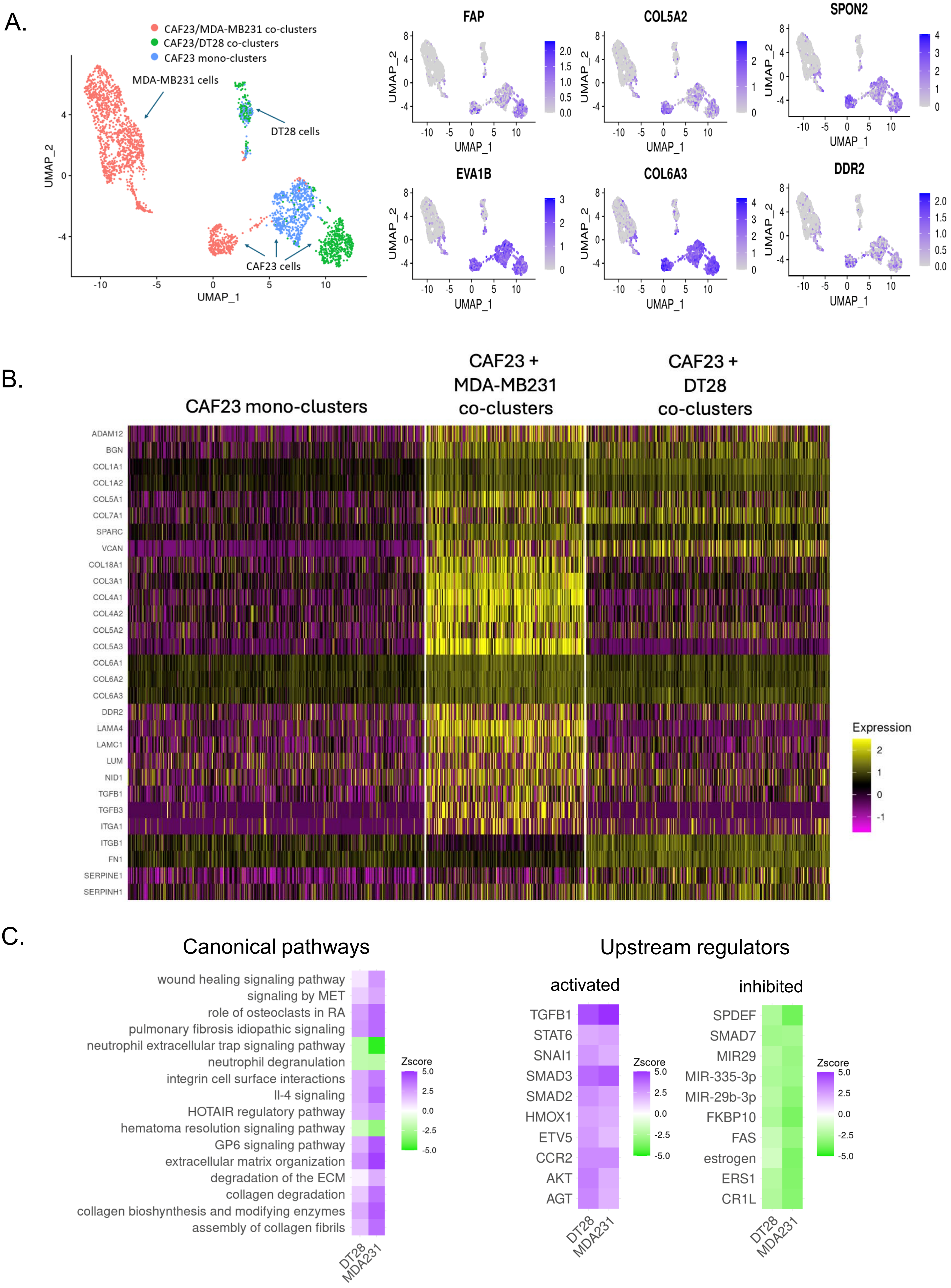

Supplemental Figure 7

A.

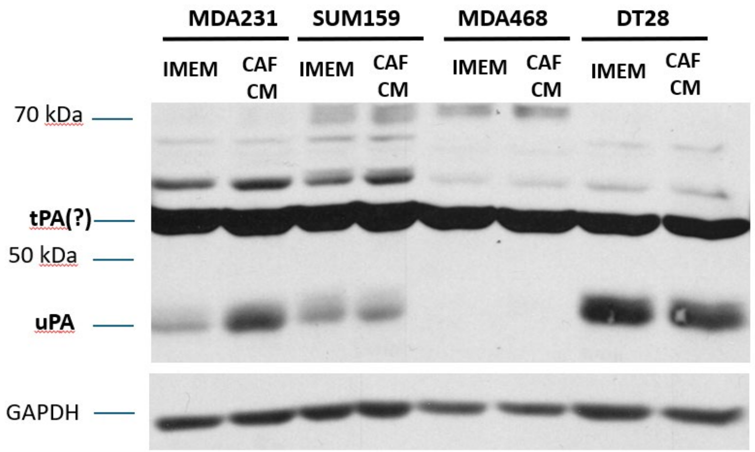


B.

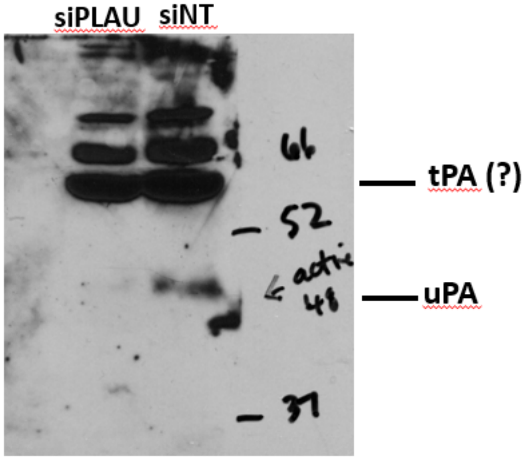


C.

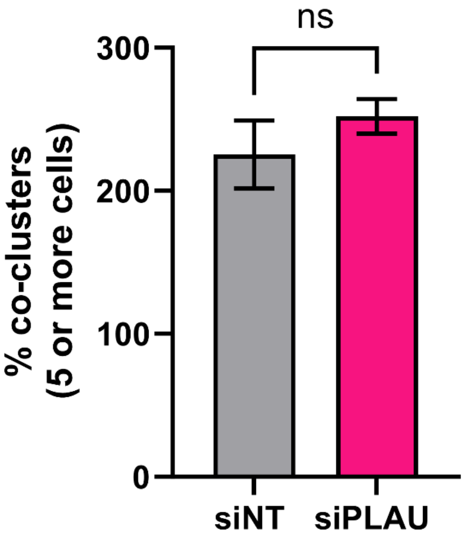

Supplemental figure 8

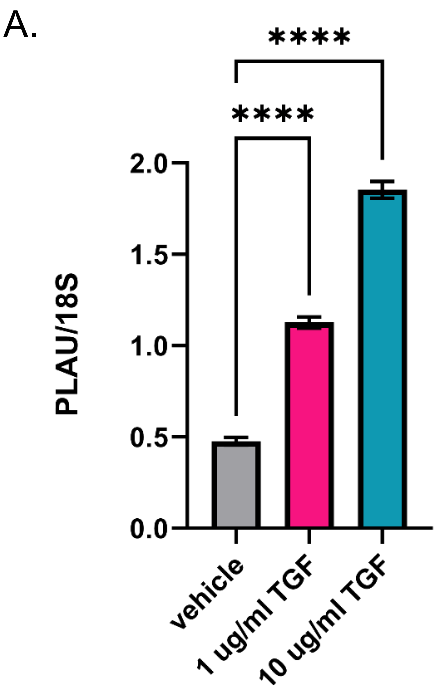


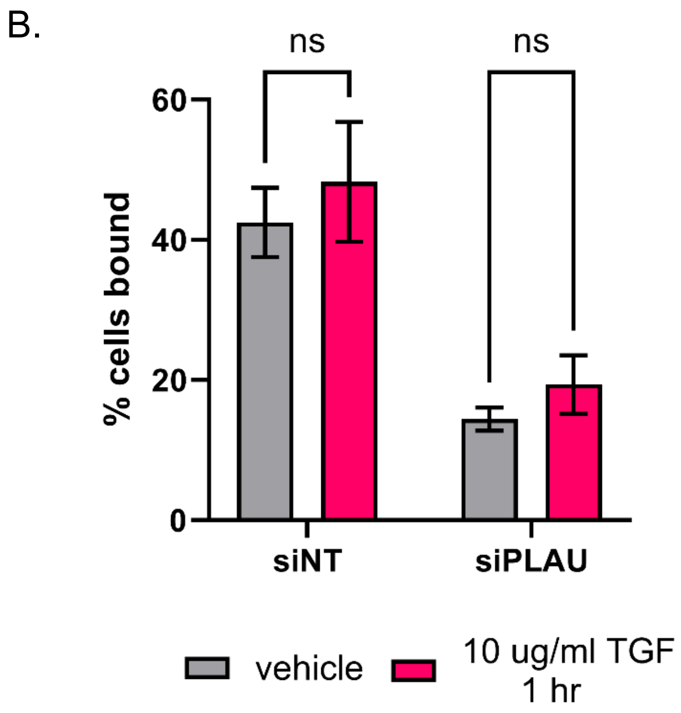


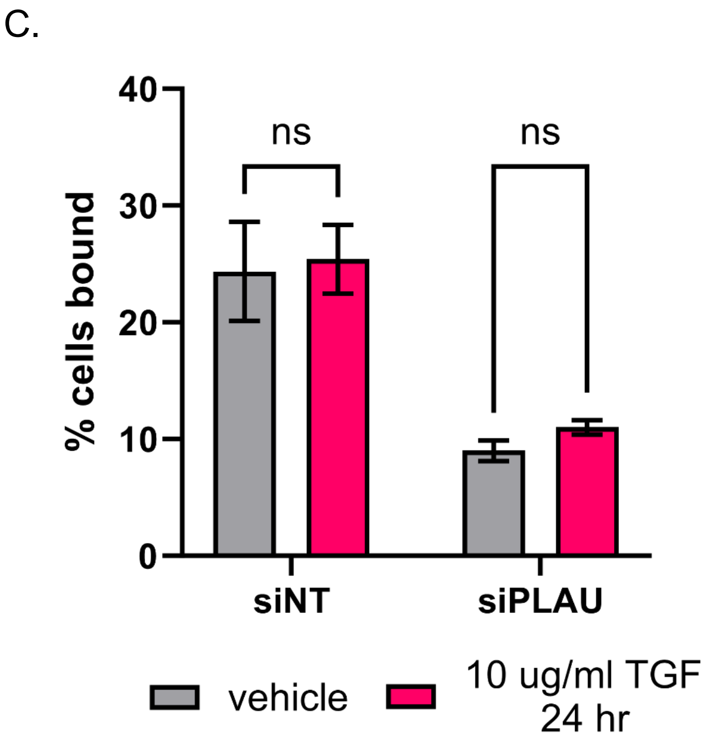
